# CO_2_-dependent opening of Connexin 43 hemichannels

**DOI:** 10.1101/2025.01.13.632711

**Authors:** Valentin-Mihai Dospinescu, Alexander Mascarenhas, Jack Butler, Sarbjit Nijjar, Kyara de Oliveira Taborda, Sean Connors, Lumei Huang, Nicholas Dale

## Abstract

Sequence and structure comparisons between alpha and beta connexins, Cx43 and Cx26, revealed that Cx43 has a motif, the carbamylation motif, that confers CO_2_-sensitivity on a subset of beta connexins. By using a fluorescent dye loading assay, whole cell patch clamp recordings and real time measurement of ATP release via GRAB_ATP_ we have demonstrated that Cx43 hemichannels open in a highly CO_2_ sensitive manner over the range 20 to 70 mmHg. Mutational analysis confirms that the equivalent residues to those in Cx26 that have been shown to be involved in mediating the effects of CO_2_ on gating of hemichannels and gap junction channels, also mediate Cx43 hemichannel gating. Our data predicts that Cx43 will be partially open and able to release ATP at resting physiological levels of PCO_2_. We have tested this prediction in acute hippocampal slices, by showing that CO_2_-dependent enhancement of synaptic transmission can be blocked by the Cx43-selective mimetic peptide Gap26. Our data resolves an inconsistency in the literature between *in vivo* studies suggesting that Cx43 hemichannels are at least partially open at rest, and *in vitro* studies, performed in the absence of HCO_3-_/CO_2_ buffering that show Cx43 hemichannels are shut. Our evidence suggests that the ancestral gene that duplicated to give the alpha and beta connexin clades must have possessed the carbamylation motif. CO_2_ sensitivity is thus a fundamental ancient characteristic of several connexins that has been lost in more recently derived members of this gene family.

## Introduction

There are 21 connexin genes encoded in the human genome. Connexins are transmembrane proteins that can form hemichannels and gap junction channels. The existence of 21 isoforms suggests functional specialisation across the gene family (Sohl & Willecke, 2004). On the basis of molecular phylogeny, the connexins have been divided into 4 broad clades - the alpha, beta, and gamma and delta groupings (Mikalsen *et al*., 2021).

Certain members of the β-clade, Cx26, Cx30 and Cx32, are directly sensitive to CO_2_ (Huckstepp *et al*., 2010; Dospinescu *et al*., 2019). The mechanism of CO_2_ sensitivity has best been studied in Cx26. While Cx26 hemichannels are opened in response to CO_2_ (Huckstepp *et al*., 2010), CO_2_ closes Cx26 gap junction channels (Nijjar *et al*., 2021). The CO_2_-dependent gating of Cx26 hemichannels and gap junction channels involves carbamylation of a specific lysine, K125, which is part of a critical carbamylation motif, _125_KVRIEGS_131_ (Meigh *et al*., 2013). This motif is essential for the CO_2_-sensitivity of Cx26, as it allows the formation of a salt bridge between the carbamylated K125 and R104 of the neighbouring subunit (Meigh *et al*., 2013). The formation of this “carbamate” bridge induces conformational changes, ultimately leading to the opening of the hemichannel or closing of the gap junction channel (Meigh *et al*., 2013; Nijjar *et al*., 2021; Brotherton *et al*., 2022; Brotherton *et al*., 2024). Mutation of K125 to arginine, an amino acid that cannot be carbamylated, or R104 to alanine so that it cannot form a carbamate bridge, removes the ability of CO_2_ to open Cx26 hemichannels (Meigh *et al*., 2013) or close Cx26 gap junction channels (Nijjar *et al*., 2021; Nijjar *et al*., 2025). Furthermore, the introduction of the carbamylation motif into a CO_2_-insensitive connexin, Cx31, results in CO_2_-dependent hemichannel opening of Cx31 (Meigh *et al*., 2013). Other members of the β-clade, Cx30 and Cx32, are also CO_2_-sensitive, but have different sensitivity profiles (Huckstepp *et al*., 2010; Dospinescu *et al*., 2019). Evolutionary comparisons show that the carbamylation motif has been conserved over 400 million years (Dospinescu *et al*., 2019) and appears to be a necessary and sufficient feature for the CO_2_-sensitivity of connexins.

Sequence comparisons of the β-connexins and α-connexins, revealed the presence of a potential carbamylation motif within the α-connexin, Cx43. Cx43 has a motif (_144_KVKMRGG_150_) homologous to the motif in Cx26 (_125_KVRIEGS_131_) suggesting that it might also be sensitive to CO_2_. In this study we show that Cx43 hemichannels are indeed directly sensitive to changes in the partial pressure of CO_2_ (PCO_2_) over the physiological concentration range. Unlike Cx26, the CO_2_-dependent opening mechanism of Cx43 is more complex, involving additional residues beyond the homologs of K125 and R104 in Cx26. Our discovery that Cx43 hemichannels are directly CO_2_ sensitive has major physiological implications and may explain why Cx43 hemichannels appear to be open under physiological conditions but closed *in vitro* where they are usually studied in the absence of CO_2_/HCO_3-_buffers.

## Results

### Identification of a potential carbamylation motif in Cx43

Cx43, an alpha-connexin, possesses a carbamylation motif homologous to that of the beta-connexins (Fig 1A). AlphaFold3 (AF3) predicts a structure very close to the experimentally determined structures for Cx43 (Lee *et al*., 2023a; Qi *et al*., 2023) but has the advantage of including portions of the structure that have not been experimentally resolved. This is a valid approach, as the *de novo* predictions by AF3 for the cytoplasmic loop of Cx26 have been supported by a recent experimentally-derived structure (Brotherton *et al*., 2024). By performing alignments of the Cx43 and Cx26 AF3 structures (Fig 1B), we found that two residues in Cx43, K144 and K105, aligned with the residues in Cx26 that are essential for hemichannel opening to CO_2_, K125 and R104 (Fig 1C). The predicted 3D orientation of the alpha-helices involved are positioned in a similar manner to allow potential interaction between K144 and K105 following carbamylation. Cx43 also presents another feature that is similar to Cx26: the high incidence of Lys residues. In Cx26, 5 lysine residues in the cytoplasmic loop can be carbamylated (Nijjar *et al*., 2025), suggesting an environment favourable for this to happen. The presence of multiple Lys residues in the cytoplasmic loop of Cx43 implies that the local environment may be conserved to facilitate the required pKa for CO_2_ binding and carbamate bridge formation.

**Figure 1:**
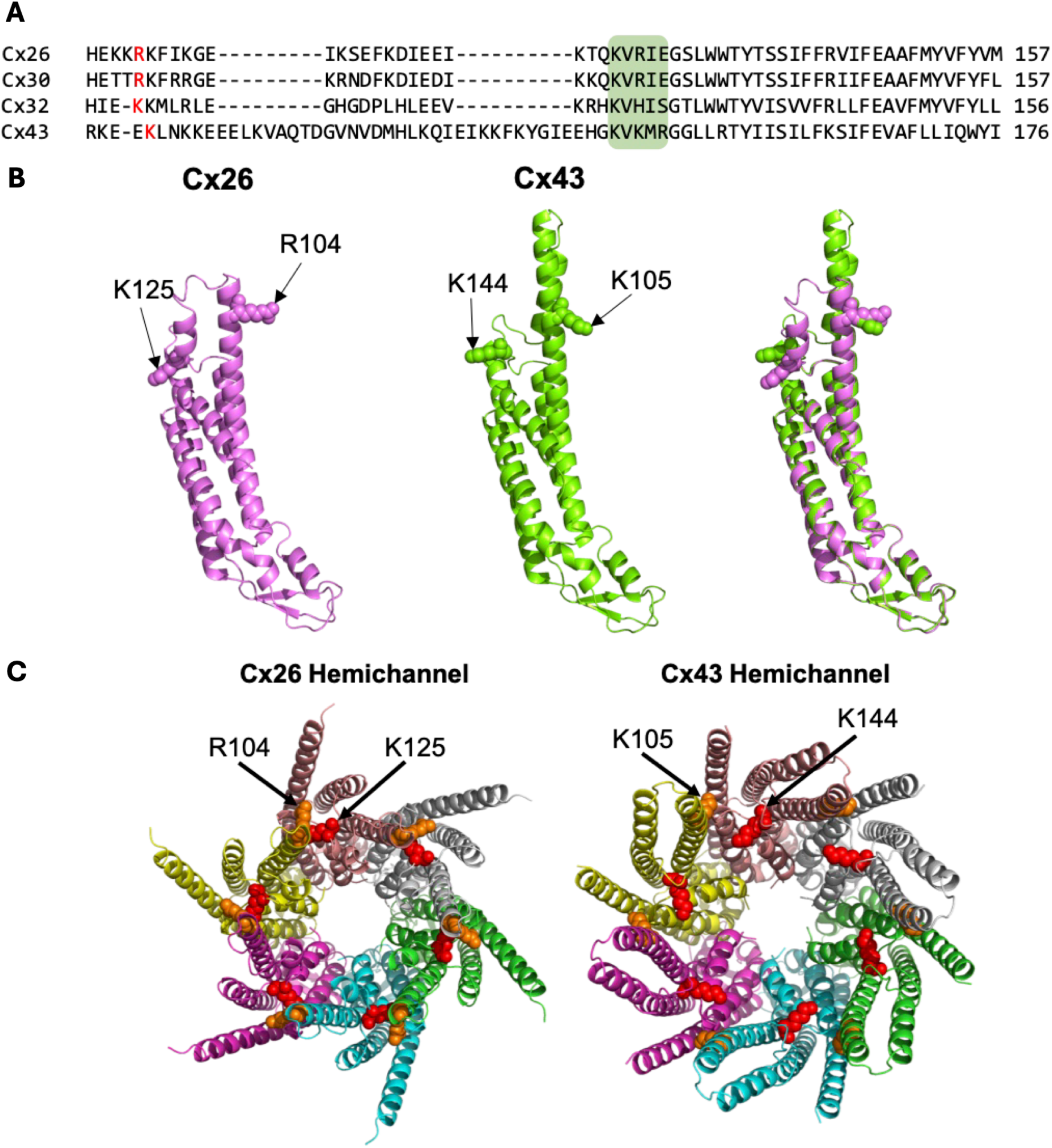
Structures of Cx43 and Cx26 predicted by AF3. (**A**) α- and β-connexin sequence alignment with the known CO_2_-sensitive connexins at the top, Cx26, Cx30 and Cx32, followed by the α-connexin Cx43. The carbamylation motif is highlighted by the green box. (**B**) Cx26 and Cx43 AlphaFold3 structural alignments. (**C**) Cx26 and Cx43 hemichannel structural prediction.

### Cx43 hemichannels are opened by CO_2_

We used an established dye-loading protocol, previously used for CO_2_ connexin studies (Huckstepp *et al*., 2010; Meigh *et al*., 2013; Meigh *et al*., 2014; de Wolf *et al*., 2016; de Wolf *et al*., 2017; Cook *et al*., 2019; Dospinescu *et al*., 2019; van de Wiel *et al*., 2020), to test whether Cx43 hemichannels are CO_2_ sensitive. We observed significant dye-loading in Cx43-expressing cells when exposed to a high CO_2_ stimulus (70 mmHg) when starting from a baseline of 20 mmHg (MW test, PCO_2_ 20 vs 70 mmHg: p = 0.008, n=5, Fig 2A). We used a zero Ca^2+^ aCSF to unblock hemichannels as a positive control to demonstrate that for WT Cx43, the zero Ca^2+^ stimulus and 70 mmHg PCO_2_ gave a similar amount of dye loading (MW test: p-value = 0.726, n=5).

**Figure 2:**
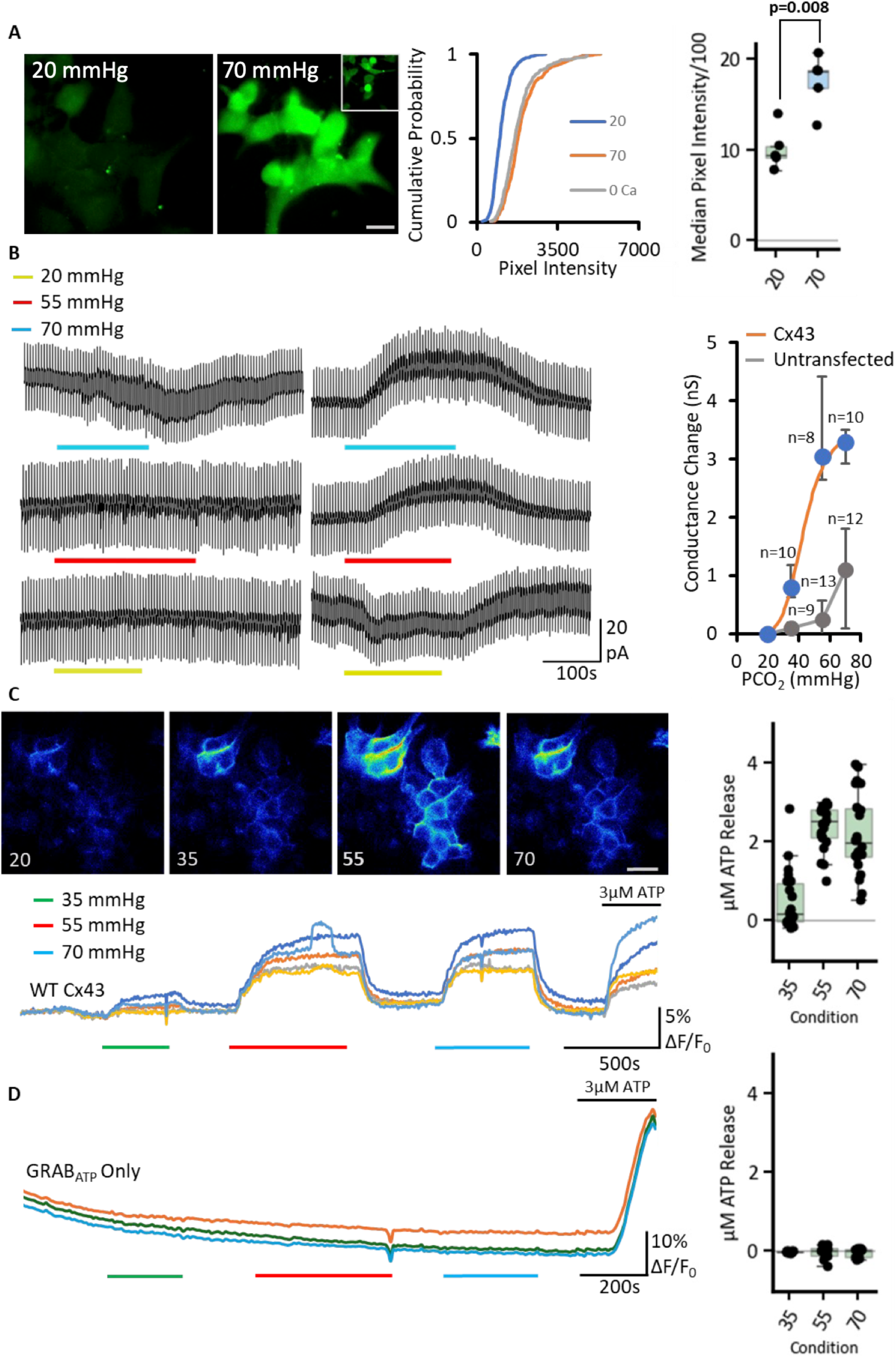
Cx43 hemichannels can be opened by an increase in PCO_2_. The CO_2_ sensitivity of Cx43 was assayed using three different methods. **(A)** Dye-loading with CBF from 20 (left) to 70 mmHg PCO_2_ (right). The inset represents the 0 Ca^2+^ control, white scale bar is 20 μm. Cumulative probability graph represents all the data points measured with more than 200 cells per condition, box plot shows the medians from each independent transfection. **(B)** Whole cell patch recordings performed on untransfected parental HeLa cells and HeLa cells expressing Cx43 at a holding potential of −50 mV with steps to −40 mV to measure whole cell conductance. The baseline was set to 35 mmHg PCO_2_ and three different concentrations were used (20, 55, 70 mmHg). The absolute conductance change evoked by a change in PCO_2_ was measured and plotted as a dose response curve (right side, median with interquartile range) with 20 mmHg being assigned to zero and the absolute value of all conductance changes plotted relative to this (see Methods for details). The points were fitted by the Hill equation H=6, EC_50_=43 mmHg (blue points, orange curve). The data for untransfected parental HeLa cells was plotted the same way (grey). **(C)** The genetically-encoded ATP sensor GRAB_ATP_ was co-transfected alongside Cx43 into HeLa cells (n=22). Representative images at different PCO_2_ concentrations of the cells are shown, grey scale bar is 20 μm. Traces from the recording the images were selected from are shown; each line represent a different cell, the fluorescence was normalised by dividing all values by the baseline median pixel intensity before plotting. Box plot shows the µM ATP release as determined by normalisation to the 3 µM control solution applied. **(D)** Parental HeLa cells transfected only with GRAB_ATP_ do not exhibit CO_2_ dependent ATP release (n=14), summary box plot shows the µM ATP release as determined by normalisation to 3 µM ATP calibration.

To further support these findings, we next used whole cell patch clamp recordings from HeLa cells expressing Cx43 (Fig 2B). We set the holding potential to −50 mV and gave regular steps to −40 mV to measure whole cell conductance. From a baseline of 35 mmHg PCO_2_, an increase in an outward conductance was observed when switching to 55 or 70 mmHg aCSF (Fig 2B). On switching to 20 mmHg aCSF, a small conductance decrease and reduction of outward current was observed (Fig 2B). Untransfected parental HeLa cells did not display changes in outward current to changes in PCO_2_. However, some parental HeLa cells exhibited a slow increase in an inward current on transfer from 35 to 70 mmHg (Fig 2B). This was quite distinct from the outward current observed in Cx43 expressing cells (Fig 2B). Overall, the patch clamp data predicted a dose response that could be fitted by the Hill equation with a Hill coefficient of 6 and an EC_50_ of 43 mmHg. This is similar to Cx26 hemichannels which also display a steep CO_2_ dose response curve with a Hill coefficient of >4 (Huckstepp *et al*., 2010; de Wolf *et al*., 2017; van de Wiel *et al*., 2020).

To assess whether the CO_2_-dependent opening of Cx43 resulted in ATP release, we co-transfected HeLa cells with the genetically encoded ATP sensor - GRAB_ATP_ (Wu *et al*., 2022). CO_2_ dependent increase in ATP release was indeed observed (n=22, Fig 2C). We observed an increase in GRAB_ATP_ fluorescence on going from 20 to 35 mmHg PCO_2_ and further to 55 mmHg PCO_2_ (Fig 2C). While ATP release at 70 mmHg was slightly less than that at 55 mmHg, it remained markedly higher than at 35 mmHg PCO_2_ (Fig 2C). By contrast, HeLa cells that were not transfected with Cx43 showed no responses to any applied CO_2_ concentration as concluded from GRAB_ATP_ experiments (n=14, Fig 2D). Overall, our results show through three independent assays that Cx43 hemichannels are opened by modest changes in PCO_2_ around the physiological norm and will be partially open at a PCO_2_ of 35 mmHg (Fig 2). Once opened by CO_2_, Cx43 hemichannels display permeability to ATP (Fig 2C).

Cx43 displays sensitivity to intracellular pH via a sensor located in the C-terminus (Ek-Vitorín *et al*., 1996). We assessed the effect of truncations of the C-terminus of Cx43 (truncated after residue 256, Cx43^1-256^) that remove this pH sensor. Both CO_2_ and high K^+^ induced ATP release from HeLa cells that expressed Cx43^1-256^ (Fig 2, figure supplement 1). We also measured the effect of changing PCO_2_ on intracellular pH (pH_i_) of HeLa cells. We found that transferring from 35 to 55 or 70 mmHg respectively induced median changes in pH_i_ of −0.02 (95% CI −0.02, −0.03) and −0.13 (CI −0.10, −0.15; Fig 2, figure supplement 2). These results show that the effects of PCO_2_ on Cx43 gating are independent of those involved in its pH sensitivity.

Gap Junction Channels (GJCs) of Cx26 are closed by CO_2_ also acting via the carbamylation motif. We therefore tested whether Cx43 GJCs might also be CO_2_ sensitive by using a dye transfer assay (Nijjar *et al*., 2021). Cx43 GJCs allowed equally fast permeation of dye at all levels of PCO_2_ and therefore, unlike the hemichannel, were insensitive to CO_2_ (Fig 2, figure supplement 3).

### Identifying the residues required for CO_2_-sensitivity of Cx43

Having established the dose-response characteristics of Cx43 to CO_2_, we next examined the potential underlying mechanisms for CO_2_-dependent gating via targeted mutagenesis of possible key residues, starting with the equivalents of those known to be involved in the CO_2_-sensitivity of Cx26 (Fig 1).

Mutation of K144 to glutamine (K144Q), an amino acid that cannot be carbamylated, partially reduced the CO_2_-sensitivity of the channel: the 70 mmHg stimulus resulted in significantly less increase in dye loading than the zero Ca^2+^ positive control (Fig 3A, MW test: p = 0.048, n=5). However, mutation of Lys105 to a non-charged residue (K105Q) resulted in hemichannels that gave a similar increase in dye loading to the zero Ca^2+^ positive control (Fig 3B, MW test: p = 0.111, n=5). In Cx26, mutation of the residues involved in carbamylation (K125 and R104) results in complete loss of CO_2_ sensitivity (Meigh *et al*., 2013; Nijjar *et al*., 2021). The lack of such a complete effect on the CO_2_ sensitivity of Cx43 implies that the mechanism is more complex and may involve further residues. Interestingly, mutation of Lys144 to Arg (K144R) completely abolished the CO_2_ sensitivity of Cx43 hemichannels as measured by dye loading (Fig 3, figure supplement 1). The difference between the effect of mutating Lys144 to Arg and Gln on CO_2_ sensitivity may arise because the substitution to Arg introduces a positive charge. Since K144 appears to be carbamylated, its primary amine must remain unprotonated so that the unpaired electrons of the nitrogen can mount a nucleophilic attack on the carbon of CO_2_ to form the carbamate bond. Thus, the K144Q mutation may preserve this charge and be a better substitute that removes the capacity for carbamylation while preserving the overall charge within this region of the protein. Consequently, we have predominantly used the Lys to Gln substitution in this study.

**Figure 3:**
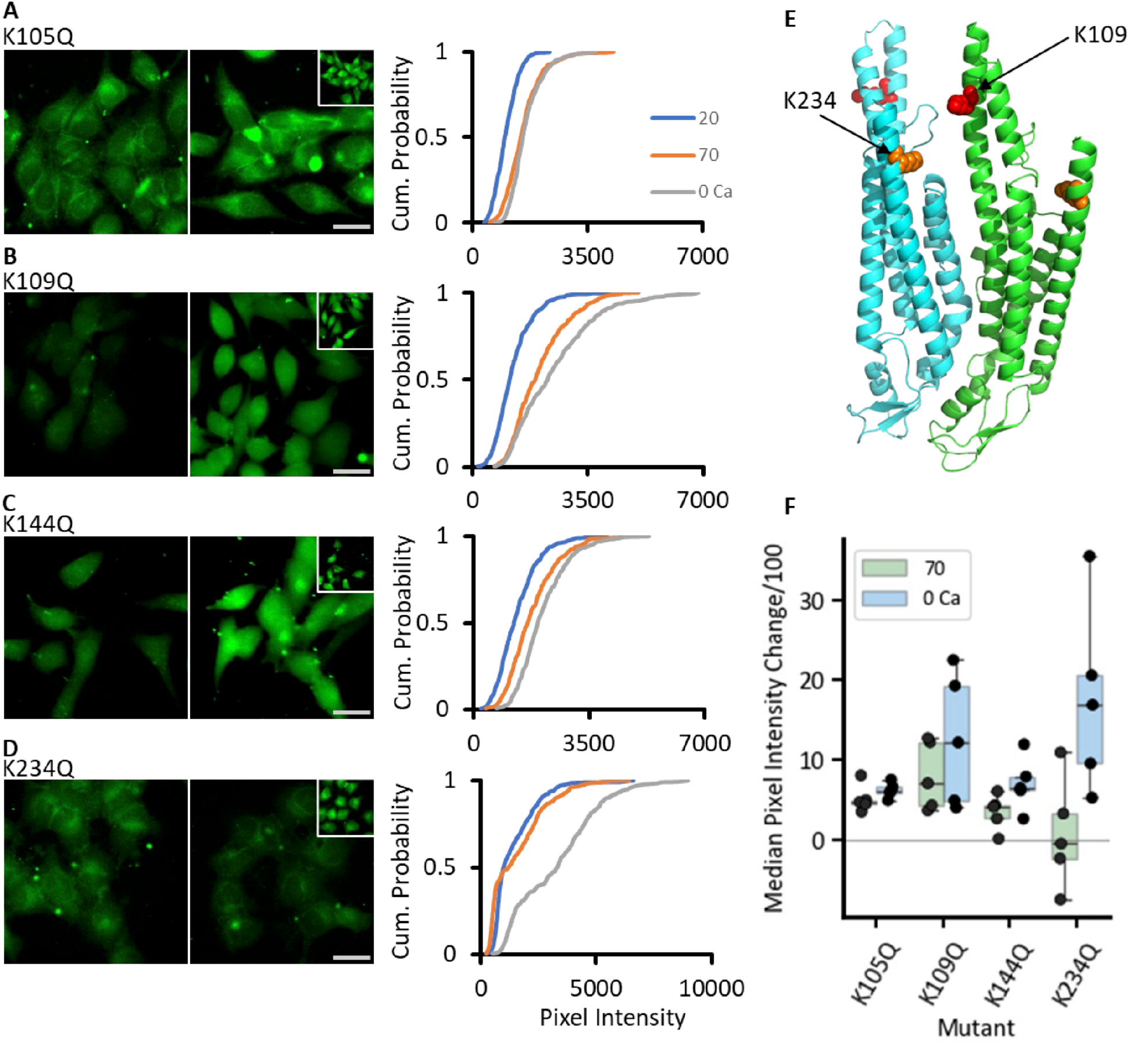
The effect of single Lys mutations on the CO_2_ sensitivity of Cx43. HeLa cells transfected with the mutant variant of Cx43 were subjected to the dye-loading protocol. Images of dye loading in response to CO_2_ and the 0 Ca^2+^ positive control (insets), together with the cumulative probability plots are shown for K105Q (**A**), K109Q (**B**), K144Q (**C**), K234Q (**D**). **(E)** AlphaFold3 prediction of Cx43 with hypothesised carbamylation bridge residues highlighted – K234 in orange and K109 in red. **(F)** Summary box plots for dye-loading showing the change in median pixel intensity from 20 mmHg PCO_2_ for each condition (70 mmHg PCO_2_ – green and 0 Ca^2+^ blue, n=5) for WT (recalculated data from Fig 2A for comparison) and each mutation. Scale bars are 20 µm.

We further inspected the AF3 structure prediction and identified two further lysine residues of interest (Fig 3E). K109 in Cx43 aligns with K108 of Cx26, a residue known to be carbamylated and required for Cx26 gap junction closure (Nijjar *et al*., 2025). AF3 also suggests that K109 could potentially interact with K234 following any carbamylation (Fig 3E). We therefore mutated both residues to Gln to test their possible involvement in CO_2_ sensitivity. The dye-loading assay showed that K109Q remained CO_2_ sensitive (Fig 3C MW test vs zero Ca^2+^ control, p = 0.274, n=5). However, the mutation K234Q displayed impaired CO_2_ sensitivity (Fig 3D, MW test vs zero Ca^2+^ control: p = 0.016, n=5).

As the dye loading assay suggested that single mutations were on the whole rather ineffective in abolishing CO_2_ sensitivity, we assayed ATP release via GRAB_ATP_ to gain further supporting evidence. We found that CO_2_-evoked ATP release remained dose dependent and very similar to the WT Cx43 in the K105Q (n=23) and K144Q (n=12) mutants (Fig 4A, C, Figure 4, figure supplement 1). Cx43 hemichannels with the K109Q mutation appeared to have their sensitivity to CO_2_ shifted to lower values of PCO_2_ (n=20), appearing to give maximal ATP release at 35 mmHg (Fig 4B,E, Fig 4 figure supplement 1). K234Q Cx43 hemichannels had greatly reduced sensitivity to CO_2_ (n=15, Fig 4D, F). To check that K234Q did not have some non-specific effect on hemichannel function, we used high K^+^ induced depolarisation as a way of opening the hemichannel independently of changes in PCO_2_ as a positive control. This stimulus resulted in robust ATP release.

**Figure 4:**
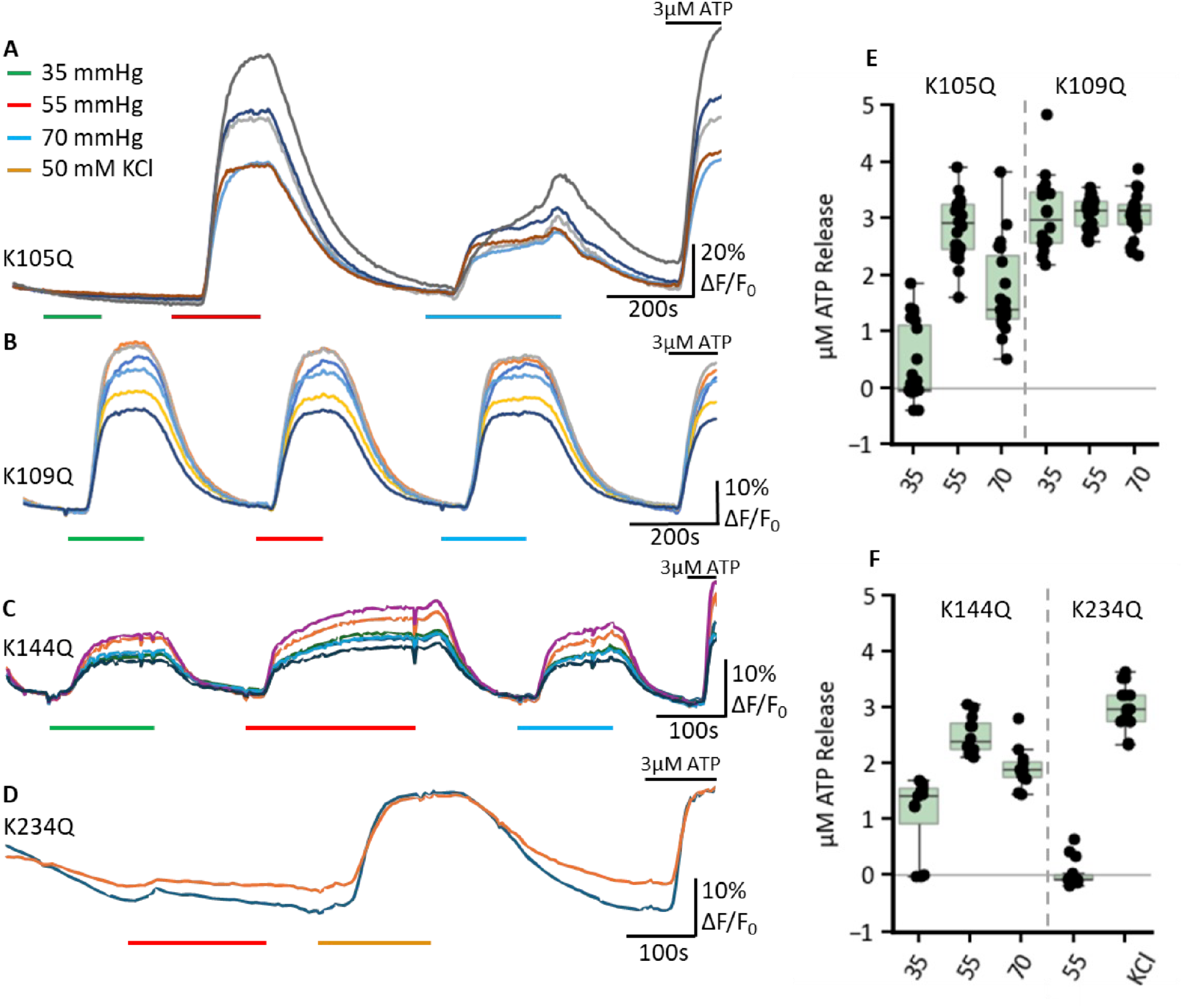
The effect of single Lys mutations on CO_2_-dependent ATP release from Cx43. The genetically-encoded ATP sensor GRAB_ATP_ was co-transfected alongside Cx43 mutant variations, K105Q (**A**) n=23, K109Q (**B**) n=20, K144Q (**C**) n=12, K234Q (**D**) n=15. Representative trace measurements were selected; each line represents a different cell; the fluorescence was normalised by dividing all values by the baseline median pixel intensity before plotting. Box plot on the right (**E**) and (**F**) represent the µM ATP released as determined through normalisation to the 3 µM ATP application.

### CO_2_ sensitivity of Cx43 involves multiple Lys residues

As single Lys mutations mostly had partial effects on the CO_2_ sensitivity of Cx43 hemichannels, we next mutated pairs of Lys residues. Using the dye loading assay we found that all combinations of paired mutations abolished CO_2_ sensitivity relative to the zero Ca^2+^ control (Fig 5): K109Q, K144Q (MW test p = 0.008, n=5); K105Q, K144Q (MW test p = 0.004, n=5); K105Q, K109Q (MW test p = 0.004, n=5); K105Q, K234Q (MW test p = 0.004, n=5); and K144Q, K234Q (MW test p = 0.004, n=5). Using the GRAB_ATP_ assay we confirmed that all of these double mutations also abolished CO_2_-dependent ATP release (MW test for all mutations, CO_2_ vs high K^+^ positive control p < 10^−6^, Fig 6).

**Figure 5:**
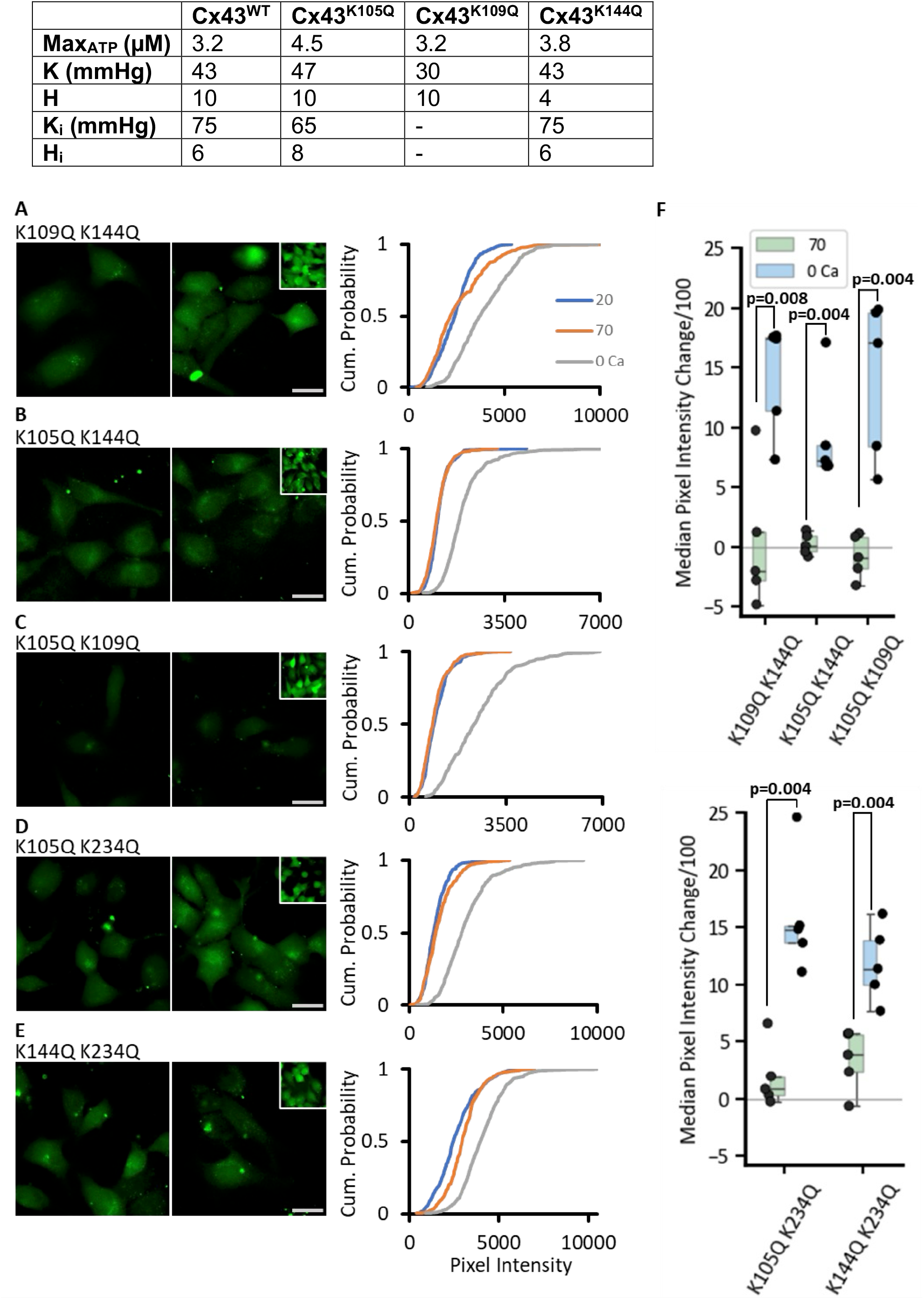
Paired Lys mutations are required to abolish the CO_2_ sensitivity of Cx43. Representative images for all combinations tried are shown – K109Q K144Q (**A**), K105Q K144Q (**B**), K105Q K109Q (**C**), K105Q K234Q (**D**), K144Q K234Q (**E**). Insets represent the 0 Ca^2+^ positive control. For each construct, the pixel intensity in expressing cells was measured from 5 individual transfections, with at least 40 cells per condition. Cumulative probability plots display all measured data points. (**F)** The box plots shows the change in median pixel intensity from 20 mmHg PCO_2_ for each transfection for 70 mmHg PCO_2_ (green) and 0 Ca^2+^ (blue).

**Figure 6:**
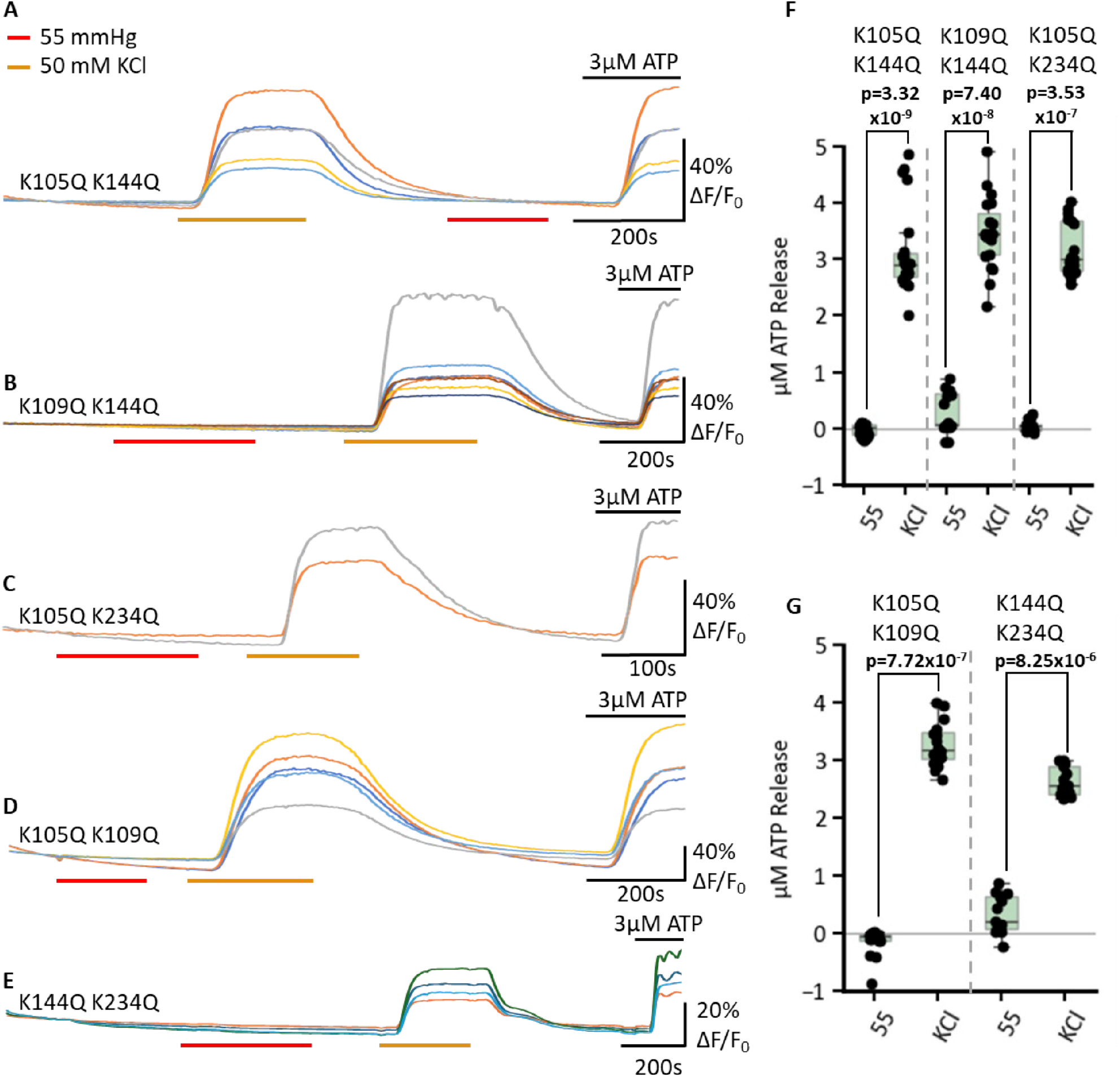
Paired Lys mutations abolish CO_2_ dependent ATP release via Cx43. HeLa cells co-expressing Cx43 and GRAB_ATP_ were subjected to changing PCO_2_-levels and fluorescence was recorded. Representative traces (**A-E**) for each of the Cx43 mutations – K105Q K144Q (**A**) n=23, K109Q K144Q (**B**) n=19, K105Q K234Q (**C**) n=17, K109Q K234Q (**D**) n=16, K144Q K234Q (**E**) n=13. The traces indicate normalised fluorescence changes (ΔF/F_0_). 3 µM ATP was applied at the end of each experiment to confirm sensor functionality. Furthermore, as a positive control, 50 mM KCl was applied to depolarise the cells and confirm channel function. (**F-G**) Box plots summarising the total ATP release in µM for each double mutant in 55 mmHg PCO_2_ and 50 mM KCl. Data points represent individual measured cells.

### The effect of introducing negatively charged residues into the carbamylation motif

We next sought to mimic carbamylation by mutating Lys to Glu. This introduces into the motif the negative charge that would occur upon carbamylation of Lys. In the dye loading assay the mutation K144E resulted in a complete loss of CO_2_ sensitivity while dye loading to zero Ca^2+^ was retained (MW test, 70 mmHg vs zero Ca^2+^: p = 0.008, n=5, Fig 7B). The mutation K105E seemed to disrupt both the CO_2_-dependent and the zero Ca^2+^ evoked dye loading (Fig 7A). K234E displayed impaired CO_2_-dependent dye loading compared to that evoked by zero Ca^2+^ (MW test, 70 mmHg vs zero Ca^2+^: p = 0.048, n=5, Fig 7C).

**Figure 7:**
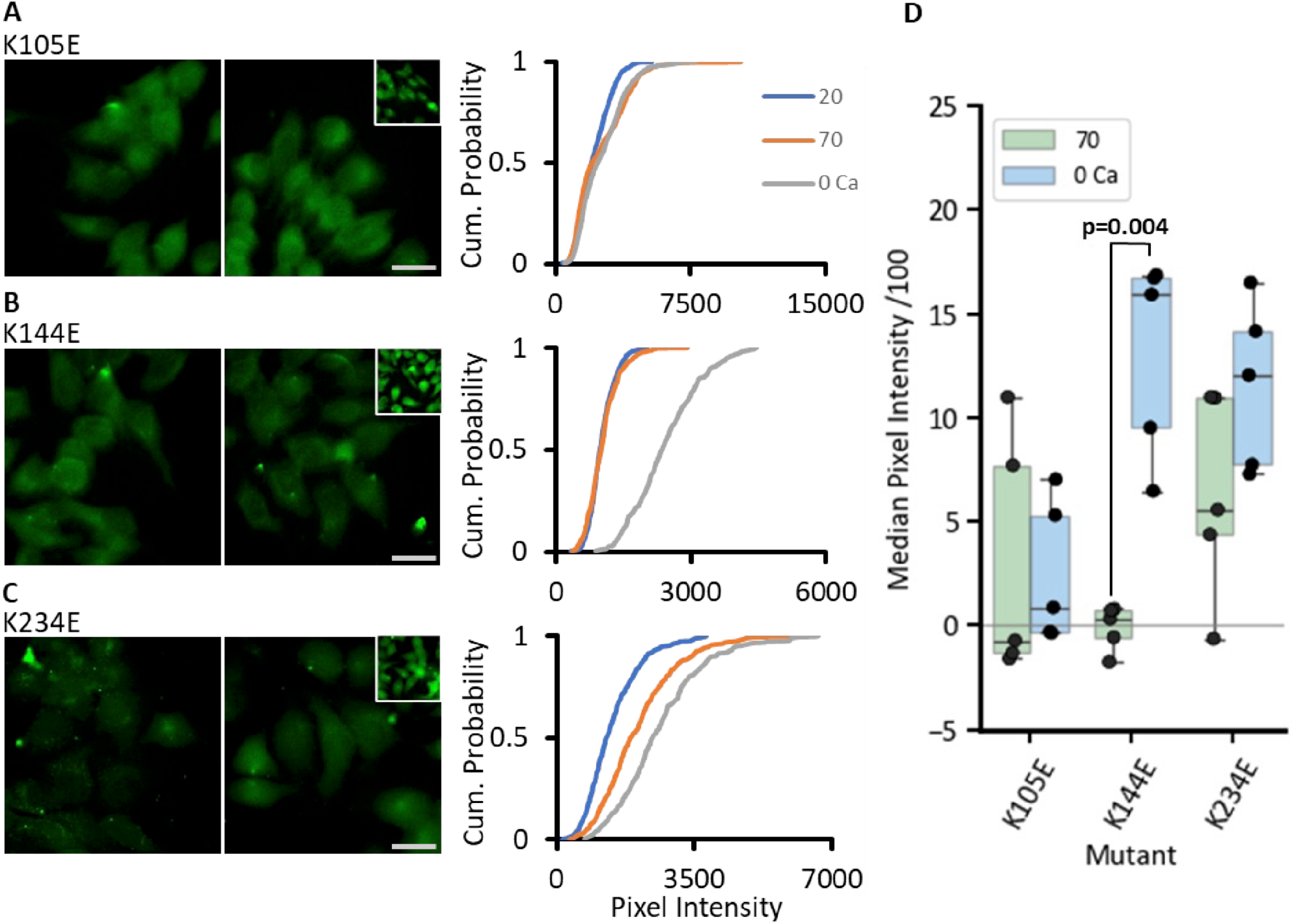
Introduction of negative charge into the carbamylation motif via Lys to Glu mutations has mixed effects on CO_2_ sensitivity of Cx43 hemichannels. Expressing cells pixel intensity for each construct was measured from 5 individual transfections, with at least 40 cells per condition. (**A-C**) Representative cell images for 20 (left), 70 (right) with insets displaying the 0 Ca^2+^ control for each mutant K105E (**A**), K144E (**B**), K234E (**C**). Scale bar represents 20 µm. Cumulative probability graphs of pixel intensities are shown on the right for each mutant with three conditions 20 mmHg PCO_2_ (blue line), 70 mmHg PCO_2_ (orange) and 0 Ca^2+^ (grey line). The box plot shows change in median pixel intensity from 20 mmHg PCO_2_ for each transfection for 70 mmHg PCO_2_ (green boxes) and 0 Ca^2+^ (blue boxes).

Using the GRAB_ATP_ assay, we found that each of these single Lys to Glu mutations disrupted CO_2_-dependent ATP release, but did not prevent depolarisation evoked ATP release (Fig 8). This disruptive effect on CO_2_ dependence might arise because introduction of the negative charge changes the local environment and makes it harder to carbamylate the other Lys residues.

**Figure 8:**
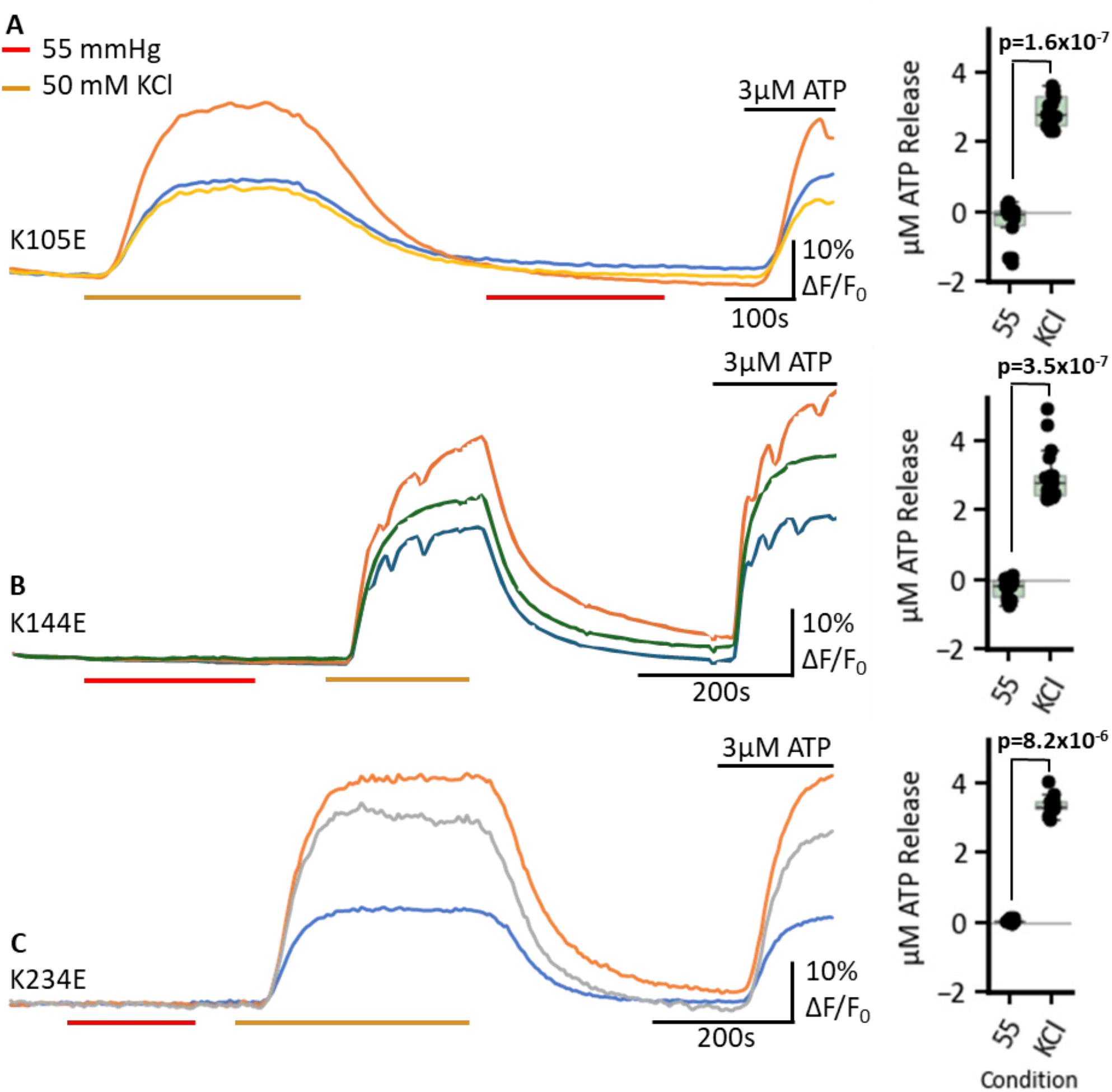
Lys to Glu mutations abolish CO_2_-dependent ATP Release. GRAB_ATP_ fluorescence traces of ATP release from cells expressing single mutant connexins: K105E (**A**) n=18, K144E (**B**) n=17, K234E (**C**) n=13, in response to 55 mmHg PCO_2_ (red bars), 50 mM KCl depolarisation control (orange bars) and a 3 µM ATP control application at the end of all experiments to confirm sensor functionality. Traces show changes in normalised fluorescence over time (ΔF/F), indicating ATP release. On the right, the box plots display the quantified ATP release in µM for each mutant under 55 mmHg PCO_2_ and 50 mM KCl. Each data point represents a measurement from an individual cell.

As a change in PCO_2_ would likely carbamylate more than one Lys residue simultaneously, we made the double mutation K105E and K109E. With the dye loading assay we found that even at a PCO_2_ of 20 mmHg, the cells loaded with dye suggesting that the mutated hemichannels were constitutively open (Fig 9A, D). Despite this cells that expressed this double mutant had normal morphology (Fig 9, movie supplement 1) and their cultures did not exhibit noticeably higher levels of cell death. Presumably the HeLa cells were able to adapt to the presence of these open channels.

**Figure 9:**
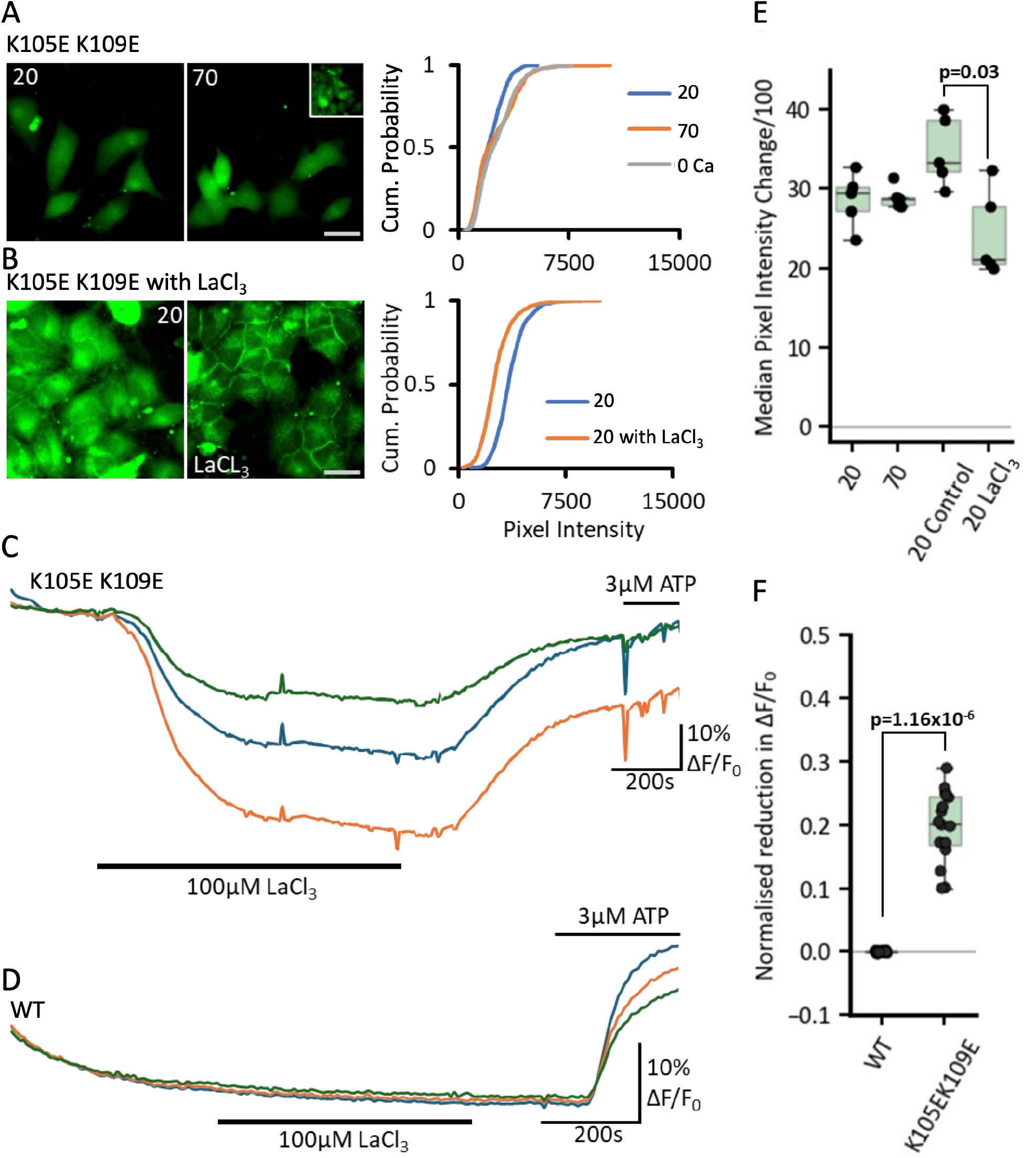
The Cx43 double mutant, K105E K109E, is constitutively open. (**A**) Representative fluorescence images of cells expressing the Cx43 K105E K109E mutant under low (20 mmHg PCO_2_, left) and high CO_2_ (70 mmHg PCO_2_, right), inset shows the 0 Ca^2+^ control. Scale bar is 20 µm. Cumulative probability plot of pixel intensity for each condition is shown on the right, overall indicating higher baseline fluorescence and perpetually open hemichannels. (**B**) Dye-loading with 20 mmHg PCO_2_ (left) and 20mmHg PCO_2_ with 100 µm LaCl_3_. Cumulative probability plot for pixel intensity under these conditions is shown on the right. (**C**) Fluorescence traces of ATP release from cells co-expressing GRAB_ATP_ and Cx43 K105E K109E (n=16) under 20 mmHg PCO_2_ and 20 mmHg PCO_2_ with 100 µM LaCl_3_ (black bar). Fluorescence is normalised to baseline. (**D**) Fluorescence traces of ATP release from cells co-expressing GRAB_ATP_ and Cx43 WT under 20 mmHg PCO_2_ and 20 mmHg PCO_2_ with 100 µM LaCl_3_ (black bar). (**E**) Box plots summarizing median pixel intensity under the different conditions, showing a significant reduction in intensity in the presence of LaCl_3_ (p = 0.03). (**F**) Box plot shows normalised fluorescence changes values for the difference between – 20 mmHg PCO_2_ (representing baseline normalisation) and the application of LaCl_3_ for both the K105E K109E and WT Cx43 constructs. **Figure 9, movie supplement 1: GRAB_ATP_ fluorescence recorded from HeLa cells expressing Cx43^K105E K109E^ during application of 100 µM LaCl_3_ in 20 mmHg PCO_2_ aCSF.**

Increasing PCO_2_ to 70 mmHg gave no further dye loading. To test whether the channels were open even at a PCO_2_ of 20 mmHg, we used the general hemichannel blocker La^3+^ (100 µM) to show that this reduced the dye loading at 20 mmHg (Fig 9B, D, MW test: p = 0.016, n=5). We used the GRAB_ATP_ assay to demonstrate that at a PCO_2_ of 20 mmHg, the K105E, K109E mutation caused continual release of ATP that could be blocked by La^3+^ (Fig 9C, D, MW test 20 mmHg vs 20 mmHg + La^3+^, p=7.7×10^−7^, n=16; Fig 9, movie supplement 1). This continual release of ATP at a PCO_2_ of 20 mmHg was not evident from cells expressing WT Cx43, as application of 100 µM La^3+^ had no effect on the GRAB_ATP_ fluorescence (Fig 9C, D).

### Physiological role for CO_2_-dependent ATP release via Cx43 from astrocytes

Cx43 hemichannels are CO_2_ sensitive and are likely to be partially open at typical levels of PCO_2_ in mammalian tissue. They could thus be able to release ATP under physiological resting conditions (35 mmHg PCO_2_, Fig 2). Considering the ubiquitous distribution of Cx43 in astrocytes (de Ceglia *et al*., 2023) and the likely resting PCO_2_ in the brain (Hogg *et al*., 1984), we postulated that, barring external modulatory influences, Cx43 hemichannels could continually release low levels of ATP in the hippocampus. Indeed, there exists compelling evidence for astrocytic Cx43 hemichannels being partially open under resting physiological conditions *in vitro* in the hippocampus (Chever *et al*., 2014). In this study when Cx43 hemichannels were blocked by the mimetic peptide, Gap26, there was a decrease synaptic strength of about 30% that was occluded by targetted knock out of astrocytic Cx43 expression (Chever *et al*., 2014).

As many investigators state that Cx43 hemichannels are shut under physiological conditions, we tested whether the results of Chever et al (2014) could be explained by the CO_2_ sensitivity of Cx43. We predicted that if PCO_2_ were lowered sufficiently to largely close Cx43, when PCO_2_ was returned to the physiological norm, Cx43 should re-open and result in an increase of synaptic strength, presumably downstream of Cx43-dependent ATP release (Chever *et al*., 2014). Because ATP can be rapidly broken down to adenosine in the extracellular space (Frenguelli *et al*., 2007; Wall & Dale, 2013), which could then act at A1 receptors to inhibit transmission and give confounding effects, we used 8-CPT (8-cyclopentyltheophylline) to selectively block A1 receptors.

We preincubated hippocampal slices in 20 mmHg PCO_2_ aCSF for 30 mins prior to recording field EPSPs (fEPSPs) in area CA1 evoked by stimulation of the Schaffer colaterals. At constant extracellular pH, in the presence of 8-CPT, elevation of PCO_2_ from 20 to 35 mmHg resulted in a 26% increase in the magnitude of the fEPSP (n=6, Fig 10A). To test whether this effect was mediated via CO_2_-dependent opening of Cx43, we used 100 μM Gap26 to see whether this blocked the increase in fEPSP induced by 35 mmHg PCO_2_. Once again the fEPSP increased by about 30% on transfer to 35 mmHg aCSF (Fig 10B), and application of Gap26 reduced the fEPSP to below its baseline value (Fig 10B, D, Friedmann 2-way ANOVA: p=0.006, n=6). This was an effect specific to the Gap26 peptide as 100 μM of the scrambled peptide did not have significant effects on the amplitude of the fEPSP (Fig 10C, D, Friedmann 2-way ANOVA: p=0.55, n=5).

**Figure 10:**
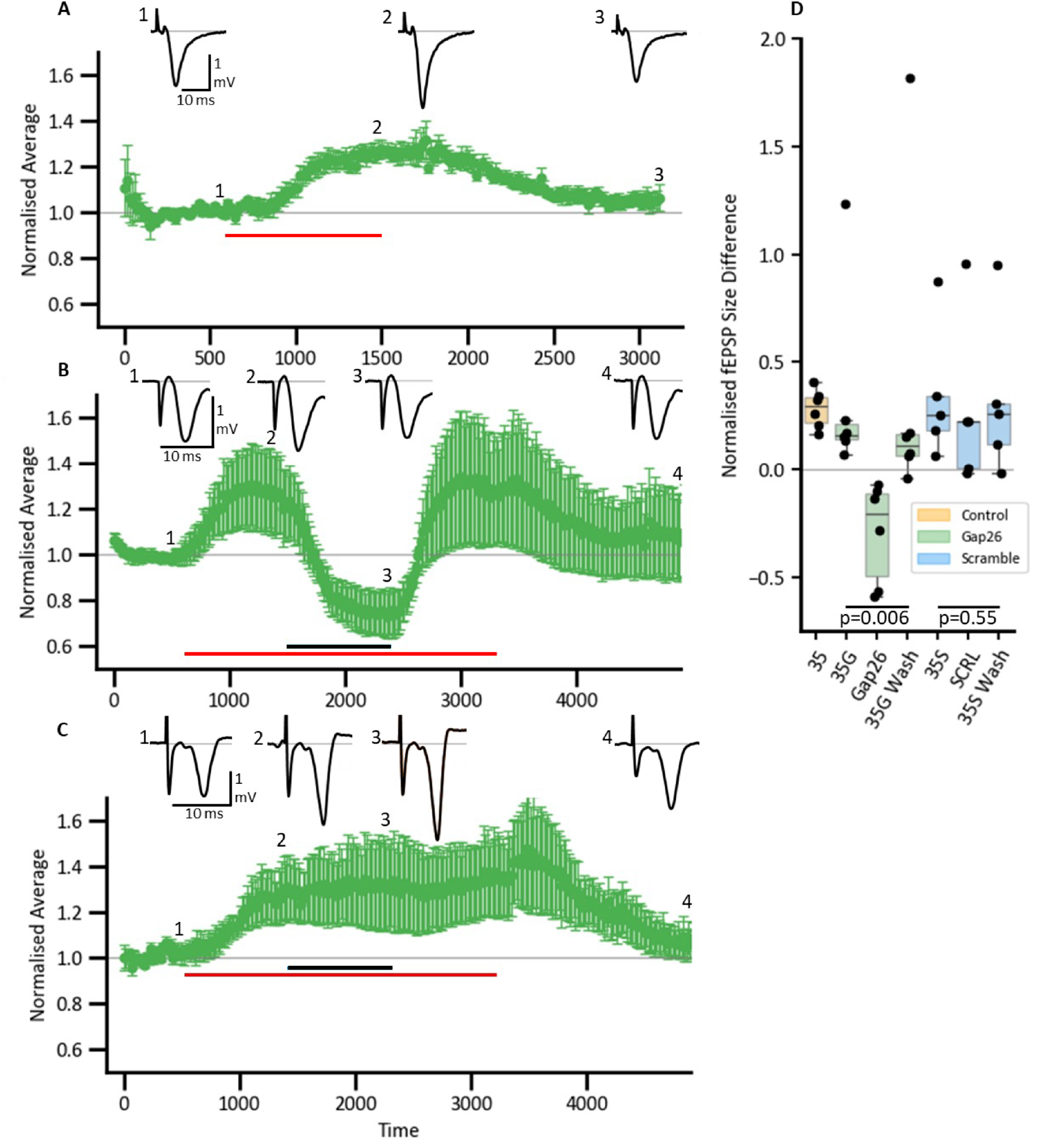
PCO_2_-dependent modulation of amplitude of fEPSPs in hippocampus is mediated via Cx43. (**A-C**) Time-course plots showing the average amplitude of the normalised fEPSP (± SEM) amplitude in response to different conditions. Inserts display representative fEPSPs. (**A**) Control condition showing an increase in fEPSP amplitude in response to a modest change to PCO_2_ (20 to 35 mmHg), red bar represent the application of 35 mmHg PCO_2_, baseline conditions – 20 mmHg PCO_2_. (**B**) EPSP responses in the presence of Gap26 (black bar) and subsequent wash. (**C**) EPSP responses with scrambled Gap26 peptide applied (black bar). (**D**) Box plots representing the normalised fEPSP size difference (baseline was subtracted) with colour coded conditions: orange – control 35 mmHg PCO_2_, green – Gap26 with pre (35G), after (35S Wash).

### Pathological mutations of Cx43 cause loss of CO_2_ sensitivity

A prominent condition associated with mutations of Cx43 is occulodental digital dysplasia (ODDD) (Laird, 2014). This syndrome involves a range of conditions such as soft tissue fusion of the digits, abnormal craniofacial bone development, small eyes and loss of tooth enamel. Later in age conditions such as glaucoma, skin disease and neuropathies may become evident. As pathological mutations of other CO_2_ sensitive connexins alter their CO_2_ sensitivity (Meigh *et al*., 2014; de Wolf *et al*., 2016; Cook *et al*., 2019; Butler & Dale, 2023), we studied two pathological mutations of Cx43. The mutation L90V causes ODDD (Shibayama *et al*., 2005; Lai *et al*., 2006) and A44V causes the skin condition, erythrokeratodermia variabilis et progressiva (EKVP) (Cocozzelli & White, 2019; Srinivas *et al*., 2019). We found that the L90V mutation removed CO_2_ sensitivity from Cx43 by both the dye loading assay and the GRAB_ATP_ assay (Fig 11). The mutation A44V blocked CO_2_ dependent ATP release (Fig 11).

**Figure 11:**
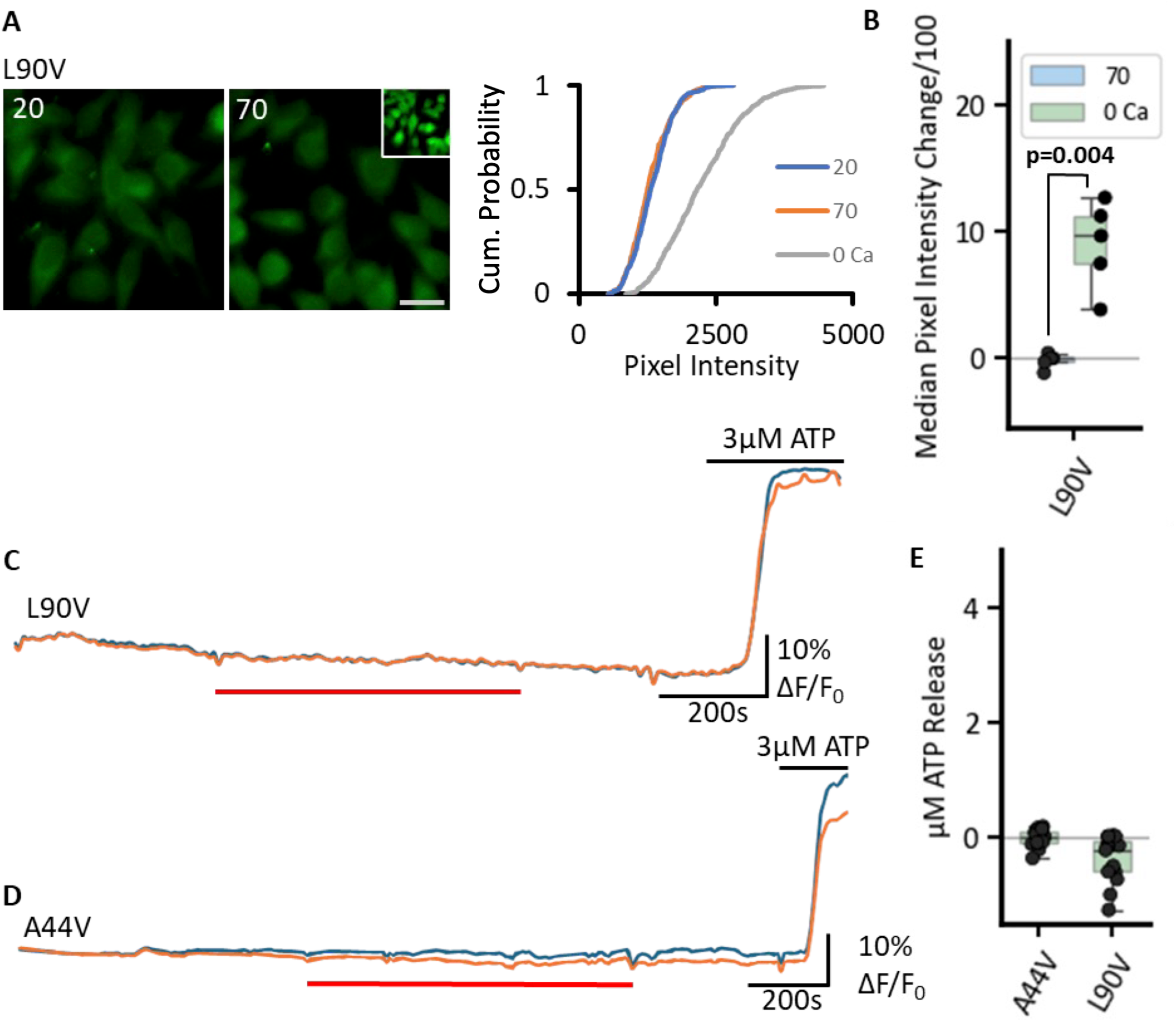
Pathological mutations of Cx43 remove its sensitivity to CO_2_. **A**,**B)** L90V prevents CO_2_-dependent dye loading. The dye loading assay shows no change in fluorescence between 20 and 70 mmHg, yet functional channels are expressed as shown by the zero Ca^2+^ positive control (inset). Box plot shows the change in median pixel intensity from 20 mmHg PCO_2_ to 70 mmHg PCO_2_ (green boxes) and 0 Ca^2+^ (blue boxes) for each transfection. **C-E)** GRAB_ATP_ recordings shown that L90V and A44V also abolish CO_2_-dependent ATP release (baseline 20 mmHg PCO_2_, red bar 70 mmHg PCO_2_).

## Discussion

### Hemichannels of Cx43 are CO_2_ sensitive

In contrast to Cx26 expression, which is restricted to specific, small regions of the brain (Nagy *et al*., 2001; Rash *et al*., 2001; Nagy *et al*., 2011), Cx43 serves as the principal astrocytic connexin, exhibiting widespread presence throughout the brain (Schulz *et al*., 2015; Yin *et al*., 2018; de Ceglia *et al*., 2023). This widespread distribution, combined with our results, suggests that Cx43 gives the capacity for CO_2_ sensitivity across the entire brain.

Our data from three independent assays (dye loading, measurement of ATP release and whole cell recordings) show that hemichannels of Cx43 open to modest changes in PCO_2_. However, Cx43 gap junction channels are insensitive to these same changes in PCO_2_. At physiological levels of PCO_2_ Cx43 hemichannels are partially open and able to release ATP. This observation potentially resolves a puzzling inconsistency in the literature. Many biophysical recordings of Cx43 hemichannels, performed in simplified buffered salines without appreciable dissolved CO_2_ or HCO_3-_, document Cx43 hemichannels as being shut (Contreras *et al*., 2003; Schalper *et al*., 2010). By contrast, under physiological conditions both *in vitro* and *in vivo*, which necessarily involve CO_2_-HCO_3-_buffering, the evidence suggests that Cx43 hemichannels remain open to some extent (Chever *et al*., 2014; Guillebaud *et al*., 2020; Turovsky *et al*., 2020). Continual ATP release from astrocytes via Cx43 has been shown to give a tone of ATP that enhances the strength of synaptic transmission in the hippocampus by about 20% (Chever *et al*., 2014). Our data is highly complementary to these observations as we find an enhancement of fEPSP amplitude of about 30% when increasing PCO_2_ from 20 to 35 mmHg. It is important to note that our experiments were performed under isohydric conditions and differ from a prior study that examined hypocapnia with simultaneous alkalinisation. This manipulation enhanced fEPSP strength through a reduction in extracellular adenosine and consequently less inhibition via A1 receptors (Dulla *et al*., 2005).

While our data suggests that Cx43 hemichannels are sensitive to PCO_2_ over the range 20-70 mmHg, there is an indication that there may be some CO_2_-dependent inhibition of hemichannel opening above 55 mmHg. Slightly less ATP release was consistently seen at a PCO_2_ of 70 mmHg compared to 55 mmHg. Furthermore, we cannot be sure that Cx43 hemichannels are completely shut at a PCO_2_ of 20 mmHg, as it is not possible to reduce the PCO_2_ further without changing to HEPES buffered salines. That Gap26 reduced synaptic transmission below the baseline amplitude recorded in slices bathed in 20 mmHg aCSF suggests that even under this condition there may still be a low level of Cx43 hemichannel gating at least in neural tissue which will continually generate CO_2_ via oxidative phosphorylation.

Cx43 gap junction channels and hemichannels are closed by intracellular acidification. This involves interaction between the C-terminus and the cytoplasmic loop (Duffy *et al*., 2002; Hirst-Jensen *et al*., 2007). Similar interactions between the C-terminus and cytoplasmic loop are involved in regulating hemichannel activity stimulated by increases in intracellular Ca^2+^ (Iyyathurai *et al*., 2018). However, the CO_2_ dependent opening of Cx43 is independent of these mechanisms as it persists when the C-terminus has been deleted.

### The residues involved in CO_2_ sensitivity and possible mechanisms

By exploiting the new structural data for connexins (Myers *et al*., 2018; Flores *et al*., 2020; Lee *et al*., 2020; Yue *et al*., 2021; Brotherton *et al*., 2022; Qi *et al*., 2022; Lee *et al*., 2023a; Lee *et al*., 2023b; Qi *et al*., 2023), the predictive power of AF3 and our understanding of carbamylation and CO_2_-dependent gating of Cx26 hemichannels and GJCs (Nijjar *et al*., 2021; Brotherton *et al*., 2024), we identified a series of residues that could plausibly be involved in mediating the CO_2_ sensitivity of Cx43: K144, the equivalent of K125 in Cx26; K105, the analogue of R104 in Cx26; K109, the equivalent of K108 in Cx26; and K234, an equivalent of R216 in Cx26. Both K125 and K108 in Cx26 are known to be carbamylated (Nijjar *et al*., 2025). Carbamylation of K125 is essential for hemichannel opening to CO_2_, whereas carbamylation of both K125 and K108 is required for GJC closure to CO_2_ (Nijjar *et al*., 2025). R216 in Cx26 has been tentatively identified as a possible interacting partner for K108 but this could be indirect (Nijjar *et al*., 2025). Crucially the spatial arrangements of K144, K105, K109 and K234 in AF3 predictions indicate that these residues are sufficiently close to interact across the subunit boundary.

In Cx26, K125 is proposed to interact with R104 following carbamylation, and interestingly mutation of either K125 or R104 is sufficient to abolish CO_2_ dependent hemichannel opening and GJC closure. Mutation of K108 is also sufficient by itself to abolish GJC closure to CO_2_ (Nijjar *et al*., 2025). By contrast single mutations to Gln of any of the 4 Lys residues identified in Cx43 as the equivalents to those involved in CO_2_-dependent gating of Cx26 were unable to completely abolish CO_2_ sensitivity. Similarly the mutations K125E and R104E in Cx26 create consitutively open hemichannels. However single Lys to Glu mutations in Cx43 did not have this effect. Our data show that in Cx43 at least 2 of the 4 identified residues must be mutated to completely abolish CO_2_ sensitivity or to give constitutively open hemichannels. For example mutation of any two Lys residues to Gln, completely abolishes CO_2_ dependent ATP release and dye loading. Conversely the double mutation, K105E and K109E, is required for constitutively open hemichannels.

This suggests that in Cx43 there is some redundancy in the effect of CO_2_. One possible interpretation is that two carbamate bridges are formed: one being between K144 and K105 (the direct equivalent of K125-R104 in Cx26); and the second being between K109 and K234. There is certainly some data that support this: the double mutations K109Q-K144Q, K105Q-K234Q and K144Q-K234Q being completely insensitive to CO_2_. These double mutations would remove both of the proposed bridges. However our observation that the mutations K105Q-K144Q, K109Q-K234Q also completely remove CO_2_ sensitivity is inconsistent with a simple two bridge hypothesis as these double mutations would only affect one of the bridges and should therefore leave the other intact to give some degree of CO_2_ sensitivity. Instead, we favour a hypothesis in which any of these four residues could be carbamylated but that they all have potential to interact. Depending on which residues become carbamylated a bridge could form between K105 and K234 or between K109 and K144, and possibly even K144 and K234. There is some support for this idea from AF3 as K105 and K109 are only one turn distant in an alpha helix and both could therefore interact with either K144 or K234 following carbamylation. The residues in Cx43 that are involved in CO_2_ sensitivity are equivalent to those identified in Cx26, where they participate in both hemichannel and gap junction channel gating to CO_2_. However in Cx43 these residues are only involved in mediating the sensitivity of the hemichannel to CO_2_. In this respect it is interesting that mutations of the different Lys residues alter the dose-sensitivity of Cx43 to CO_2_ in different ways (Fig 4, figure supplement 1). New cryo-EM studies of Cx43 hemichannels at vitrified different levels of PCO_2_ would shed more light on the mechanism of opening,

### Physiological implications of the CO_2_ sensitivity of Cx43

Cx43 is the most widely expressed connexin in the human body, being present in every organ system (Lucaciu *et al*., 2023). In metabolically highly active organs such as liver, kidney and brain, our data suggest that the physiological PCO_2_ will be sufficient to substantially open Cx43 hemichannels (Hogg *et al*., 1984). In the context of the brain, Cx43 is the main astrocytic connexin and is expressed in all subtypes of astrocytes (de Ceglia *et al*., 2023). This would imply that CO_2_-sensitivity mediated via Cx43 will extend to potentially all brain regions. As astrocytes are non-excitable, they are unlikely to become sufficiently depolarised for Cx43 hemichannels to open via voltage dependent gating. We suggest therefore, in non-excitable cells such as astrocytes, that variations in PCO_2_ may be the most important physiological regulator of Cx43 hemichannel gating. For astrocytes, CO_2_-dependent opening of Cx43 hemichannels is likely to result in the release of a mix of small, neurochemically significant molecules such as ATP, glutamate, D-serine and lactate, depending on their intracellular concentration.

Cx43 is also highly expressed in the heart, if a proportion of the population were to be in hemichannel form as opposed to GJCs (De Smet *et al*., 2021), the newly-discovered CO_2_ sensitivity of hemichannels may have profound implications for heart function given that cardiomyocytes are highly active, and will thus generate high levels of CO_2_ through oxidative phosphorylation. There is a long-established link between CO_2_ and cardiac function with effects such as increased coronary blood flow in response to CO_2_ as well as changes in myocardial contractile function (Crystal, 2015). CO_2_ can even cause arrythmias (Zhang *et al*., 2019). The effects of CO_2_ may be mediated through changes in intracellular pH, which is well known to alter the function of cardiac myocytes (Spitzer *et al*., 2002; Vaughan-Jones *et al*., 2009; Orlowski *et al*., 2025). However, given that Cx43 is directly sensitive to CO_2_, an investigation as to whether CO_2_ also has direct effects on cardiac function that are independent of consequent changes in intracellular pH seems warranted.

NF-kappa B and innate immunity exhibit sensitivity to CO_2_ (Cummins *et al*., 2010; Keogh *et al*., 2017). Macrophages are CO_2_-sensitive, although at least some of this is indirect and via pH changes mediated by carbonic anhydrase activity (Strowitzki *et al*., 2022). However, a direct effect of CO_2_ on components of the immune system cannot be excluded. In cell culture, when carbonic anhydrase activity is blocked in THP-1 monocytes, the effects of CO_2_ are significantly diminished but not abolished, suggesting the existence of a pH-independent pathway (Strowitzki *et al*., 2022). As Cx43 is expressed in monocytes it is a plausible candidate to mediate direct CO_2_ sensitivity in these cells. Furthermore, Cx43 function in macrophages is deemed critical for various physiological and pathophysiological processes (Rodjakovic *et al*., 2021). The role of CO_2_ on such pathways remains to be explored.

It is interesting that pathological mutations of Cx43 abrogate CO_2_ sensitivity. This follows a pattern seen for Cx26, where a range of mutations linked to syndromic and non-syndromic hearing loss (Meigh *et al*., 2014; de Wolf *et al*., 2016; Cook *et al*., 2019), and for Cx32, where several mutations that cause X-linked Charcot Marie Tooth Disease (Butler & Dale, 2023) alter or completely block CO_2_ sensitivity. Further studies are needed to see whether loss of CO_2_ sensitivity from Cx43 does indeed contribute to pathology.

### Evolution of CO_2_ sensitivity in connexins

Our previous findings showed that a common carbamylation motif is prevalent across the beta-connexin clade and confers CO_2_-sensitivity on the connexins that possess it (Dospinescu *et al*., 2019). This motif for example is present in shark Cx32 (which is CO_2_ sensitive). Humans and sharks last shared a common ancestor about 400-450 MYA. Our discovery that a very similar carbamylation motif is present in Cx43 and confers CO_2_ sensitivity onto their hemichannels. Examining the sequences of other human and non-human alpha connexins reveals that the carbamylation motif with appropriately oriented residues is present in human Cx50, Cx59 and Cx62 as well as Cx40, Cx46 and Cx57 of non-human species (Fig 12, Fig 12 Table Supplement). We have shown Cx50 hemichannels are CO_2_ sensitive and this also depends on the carbamylation motif (Lovatt *et al*., 2025). These findings suggest that the carbamylation motif and CO_2_ sensitivity must have been present in the ancestral connexin gene that predated the establishment of the alpha and beta connexin clades. Further work is needed to understand the evolution of the carbamylation motif and CO_2_ sensitivity, but our observations indicate that it is an ancient feature within the molecular phylogeny of the connexins, rather than being a more recently derived feature restricted to the beta connexins.

**Figure 12:**
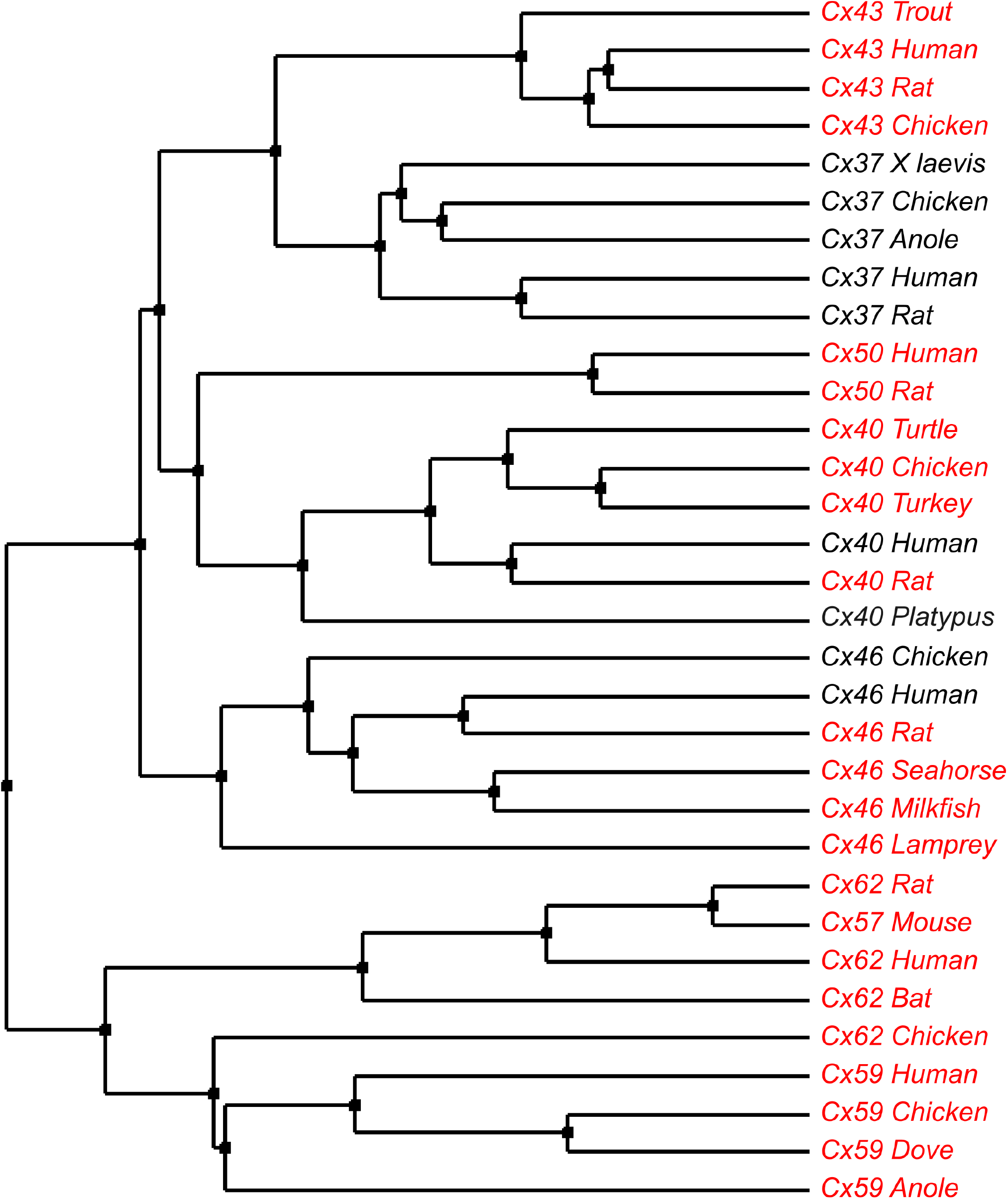
Occurrence of the carbamylation motif in the alpha connexin clade. Names in red indicate the presence of the motif. The sequences were aligned in TCoffee to check for the presence of the motif. The molecular phylogenetic tree was constructed from the aligned traces by the average distance using Blosum62 in JalView (Waterhouse *et al*., 2009). Accession numbers for the protein sequences in the tree are given in the accompanying table supplement.

## Methods

### Recording Solutions

Hypocapnic (20 mmHg PCO_2_): 140 mM NaCl, 10 mM NaHCO_3_, 1.25 mM NaH_2_PO_4_, 3mM KCl, 1 mM MgSO_4_, 10 mM D-glucose and 2 mM CaCl_2_. This was bubbled with a mix of 95%O_2_/5%CO_2_ and balanced with sufficient pure O_2_ from an oxygen concentrator to give a final pH of ~7.3.

Control (35 mmHg PCO_2_): 124 mM NaCl, 26 mM NaHCO_3_, 1.25 mM NaH_2_PO_4_, 3 mM KCl, 10 mM D-glucose, 1 mM MgSO_4_, 2 mM CaCl_2_. This was bubbled with 95%O_2_/5% CO_2_ and had a final pH of ~7.3.

Hypercapnic (55 mmHg PCO_2_): 100 mM NaCl, 50 mM NaHCO_3_, 1.25 mM NaH_2_PO_4_, 3 mM KCl, 10 mM D-glucose, 1 mM MgSO_4_, 2 mM CaCl_2_. This was bubbled with sufficient CO_2_ (~9%, balance O_2_) to give a final pH of ~7.3.

Hypercapnic (70 mmHg PCO_2_): 73 mM NaCl, 80 mM NaHCO_3_, 1.25 mM NaH_2_PO_4_, 3 mM KCl, 10 mM D-glucose, 1 mM MgSO_4_, 2 mM CaCl_2_. This was bubbled with sufficient CO_2_ (approximately 12%, balance O_2_) to give a final pH of ~7.3.

Zero Ca^2+^: 140 mM NaCl, 10 mM NaHCO_3_, 1.25 mM NaH_2_PO_4_, 3mM KCl, 1 mM MgSO_4_. On the day of recording, 10 mM D-glucose, 1 mM EGTA (Ethyleneglycol-bis(β-aminoethyl)-N,N,N',N'-tetraacetic acid) and 2 mM MgCl_2_ was added, and the solution bubbled with 95%O_2_/5%CO_2_ and balanced with sufficient pure O_2_ from an oxygen concentrator to give a final pH of ~7.3.

### Cloning and mutagenesis

Plasmids containing mutated versions of Cx43 were generated using the Gibson Assembly method (Gibson *et al*., 2009). Overlapping fragments both containing the desired mutation were PCR amplified with primers. Double mutations were cloned using successive Gibson assemblies. PCR fragments were amplified using Q5® High-Fidelity DNA Polymerase (New England Biolabs) (normal PCR or site-directed mutagenesis). The overlap region generated through PCR for Gibson Assembly was 20 base pairs, and the pCAG vector was used as the backbone. All Cx43 constructs were inserted upstream of an mCherry tag, linked via a 12 AA linker (GVPRARDPPVAT). All PCR primers were purchased from Integrated DNA Technologies (IDT). For double mutations, an additional round of cloning was performed using a single mutant as the template, to introduce the additional mutation. For the truncation (Cx43^1-256^), we created a PCR product of the truncated version of Cx43 and fused it to the same AA linker and mCherry as previously stated, to allow visualisation of cellular localisation. All constructs were confirmed by DNA sequencing (Eurofins GATC sequencing or Plasmidasaurus).

### Cell culture

HeLa DH cells were grown in Dulbecco’s Modified Eagle Medium (DMEM), supplemented with 10% fetal bovine serum, 50 μg/mL penicillin/streptomycin. HeLa DH cells were used for patch clamp studies on Cx43 and for the dye loading of all Cx43 variants. For dye loading experiments, cells were seeded onto coverslips at a density of 7.5 × 10^4^ cells per well and transiently transfected with the Cx43 constructs following the PEI Transfection Reagent protocol.

### Dye-loading

Cell expressing the construct (48-72h) were given an initial 5-minute wash with baseline solution (artificial cerebrospinal fluid – aCSF, 20 mmHg PCO_2_). Following this, the cells were exposed to either control solution, hypercapnic (70 mmHg PCO_2_), or a zero Ca^2+^ positive control (20 mmHg PCO_2_) solution, all of which contained 200 µM 5(6)-carboxyfluorescein (CBF) for 10 min. Next, cells were put into 20 mmHg PCO_2_ solution with 200 µM CBF for 5 minutes to close any open channels and prevent dye loss in the next step. Then the coverslip was washed with control solution (without CBF) for another 30 min to remove extracellular dye. For each condition a different coverslip with HeLa cells was used. Cx43 was tagged with mCherry on the C-terminus using a short linker. Expression was verified using mCherry fluorescence. The experiments were replicated using independent transfections 5 times.

Following dye loading, HeLa cells were imaged by epifluorescence (Scientifica Slice Scope, Cairn Research OptoLED illumination, 60x water Olympus immersion objective, NA 1.0, Hamamatsu ImagEM EM-CCD camera, Metafluor software). CBFwas excited by a 470 nm LED, with emission captured between 504-543 nm. Cx43 constructs had a C-terminal mCherry tag, which was excited by a 535 nm LED and emission captured between 570-640 nm.

Following acquisition analysis was performed by a person blind to the experimental condition. ImageJ (Wayne Rasband, National Institutes of Health, USA) was used to measure the extent of dye loading by drawing a region of interest (ROI) around each cell, and subsequently, the mean pixel intensity of the ROI was determined. The mean pixel intensity of a representative background ROI for each image was subtracted from each cell measurement from the same image. At least 40 cells were measured for each condition per experiment, and five repetitions of independently transfected HeLa cells were completed. The mean pixel intensities were plotted as cumulative probability distributions, and these graphs show every data point measured. To assess the effect of the 70 mmHg and zero Ca^2+^ solutions on dye loading, the difference in the median pixel intensities between the 70mmHg and 20 mmHg conditions, and the zero Ca^2+^ and 20 mmHg conditions was calculated for each transfection (Figs 3, 5, 7 and 11).

### GRAB_ATP_ recordings

Cells were transiently transfected with the pCAG-Cx43-mCherry construct and pDisplay-GRAB_ATP1.0-IRES-mCherry-CAAX (Addgene plasmid # 167582; RRID:Addgene_167582) (Wu *et al*., 2022) 48 hours prior to imaging. Cells were perfused with control aCSF until a stable baseline was reached, before perfusion with either hypercapnic or high K^+^ aCSF (positive control). Once a stable baseline was reached after solution change, cells were again perfused with control aCSF and when a stable baseline reached, recordings were calibrated by direct application of 3 μM of the corresponding analyte.

All cells were imaged by epifluorescence as above. The cpGFP in GRAB_ATP_ was excited by a 470 nm LED, with emission captured between 504-543 nm. As the Cx43 constructs were mCherry tagged, we selected only cells that expressed both cpGFP and mCherry for recording. GRAB_ATP_ fluorescence images were acquired every 4 seconds. For each condition, at least 3 independent transfections were performed with at least 2 coverslips per transfection.

Analysis of all experiments was carried out in ImageJ. Images were opened as a stack and stabilised. ROIs were drawn around cells co-expressing both sensor and connexin. Median pixel intensity was plotted as normalised fluorescence change (ΔF/F_0_) over time to give traces of fluorescence change. Amount of ATP release was quantified as concentration by normalising to the ΔF/F_0_ caused by application of 3 μM ATP.

### Intracellular pH measurement

BCECF-AM dissolved in DMSO and Pluronic f127 and diluted in 35 mmHg aCSF for a final concentration of 2 μM. Coverslips seeded with parental HeLa DH cells were incubated in BCECF for 20 minutes, before being washed in 35 mmHg aCSF for 10 minutes. Cells were perfused with control aCSF until a stable baseline was reached, before perfusion with either 55 mmHg or 70 mmHg aCSF. Once a stable baseline was reached after solution change, cells were again perfused with control aCSF. pH_i_ was then calibrated following the method in (James-Kracke, 1992). All cells were imaged by epifluorescence with the BCECF excited by a 470 nm LED and emission captured between 504-543 nm. Measurements were obtained from 3 independent transfections were performed with at least 2 coverslips per transfection.

### Patch clamp recordings

Coverslips containing non-confluent HeLa cells were placed into a perfusion chamber at room temperature and superfused with control aCSF. An Axopatch 200B amplifier was used to make whole-cell recordings from single HeLa cells. The intracellular fluid in the patch pipettes contained: K-gluconate 60 mM, Cs-gluconate 50 mM, CsCl 10 mM, TEACl 10 mM, EGTA 10 mM, Na_2_ATP 3mM, MgCl_2_ 3 mM, CaCl_2_ 1 mM, HEPES 10 mM, sterile filtered, pH adjusted to 7.2 with KOH. An agarose salt bridge was used to avoid solution changes altering the potential of the Ag/AgCl reference electrode. All whole-cell recordings were performed at a holding potential of −50 mV. Whole-cell conductance was measured by repeated steps to −40 mV. To allow for any drift in whole-cell conductance unrelated to the CO_2_ stimulus, the maximal conductance during the CO_2_ test stimulus was compared to the average of the conductance before the stimulus and after the stimulus had been fully washed off. Each whole cell recording was considered to be an independent statistical replicate.

Control (35 mmHg) aCSF was used as the baseline and switched to 20 mmHg, 55 mmHg or 70 mmHg aCSF to measure the conductance responses to differing levels of PCO_2_. To convert these responses to a PCO_2_ dose response curve, assigned a value of zero to 20 mmHg and plotted all changes relative to this. Thus, the values at 35 mmHg were the absolute values of the changes evoked by going from 35 to 20 mmHg, and the absolute value of median change from 35 to 20 mmHg was added to the changes observed going from 35 to 55 and 35 to 70 mmHg.

### Imaging assay of gap junction transfer

2-Deoxy-2-[(7-nitro-2,1,3-benzoxadiazol-4-yl)amino]-D-glucose, NBDG, was included at 200 µM in the patch recording fluid, which contained: K-gluconate 130 mM; KCl 10 mM; EGTA 5 mM; CaCl_2_ 2 mM, HEPES 10 mM, pH was adjusted to 7.3 with KOH to give a resulting final osmolarity of 295 mOsm. Cells were imaged on a Cleverscope (MCI Neuroscience) with a Photometrics Prime camera under the control of Micromanager 1.4 software. LED illumination (Cairn Research) and an image splitter (Optosplit, Cairn Research) allowed simultaneous imaging of the mCherry-tagged Cx43 subunits and the diffusion of the NBDG into and between cells. Coupled cells for intercellular dye transfer experiments were initially selected based on tagged Cx43 protein expression and the presence of a gap junctional plaque (Figure 4, figure supplement 1). Dye permeation between cells was measured at PCO_2_ levels of 20, 55 and 70 mmHg. After establishing the whole cell mode of recording, images were collected every 10 seconds. The assay was performed as described by Nijjar et al (Nijjar *et al*., 2021). The start of recording was taken to be the first image following the establishment of a stable whole cell recording. For each gap junction recording (considered as an independent statistical replicate), analysis of the cell images was performed in ImageJ using an ROI drawn on each cell to measure the median pixel intensity of the donor and acceptor cells. The time for the median fluorescence of the acceptor cell to reach 10% of the donor cell was then determined and used as the value for statistical comparisons.

### Hippocampal slice preparation

Mice of either sex, 3-5 weeks old, were killed by cervical dislocation and decapitated in accordance with United Kingdom Animals (Scientific Procedures) Act (1986). The brain was dissected and kept on ice both the cerebellum and the rostral section of the brain were removed. 400 µm thick sagittal slices were cut with a vibratome (Microm HM 650V microsliver) in ice-cold cutting solution composed of (in mM: 85 NaCl, 2.5 KCl, 0.5 CaCl_2_, 1.25 NaH_2_PO_4_, 24 NaHCO_3_, 25 glucose, and 75 sucrose; pH was adjusted to 7.4, bubbled with 95% O_2_ + 5% CO_2_). Then slices were incubated in aCSF composed of (in mM: 127 NaCl, 1.9 KCl, 2 MgCl_2_, 2 CaCl_2_, 1.2 KH_2_PO_4_, 26 NaHCO_3_, 10 D-glucose, pH 7.4, when bubbled with 95% O_2_ and 5% CO_2_) at 33 °C. For fEPSP recordings an individual slice was pre-incubated in 20 mmHg PCO_2_ aCSF for 30 mins, following which the slice was transferred to a recording chamber submerged in aCSF at 32°C. For stimulation, pulses were delivered by a stimulator via concentric bipolar metal stimulating electrode placed on the surface of CA3. For extracellular recordings, an aCSF-filled microelectrode was placed in CA1.

### Statistics

Data is plotted as either cumulative probabilities (showing every data point) or box and whisker plots where the box is interquartile rage, bar is median, and whisker extends to most extreme data point that is no more than 1.5 times the interquartile range. All individual data points are superimposed on the plots.

In the case of dye loading one transfection was considered one replicate. In the case of patch clamping and GRAB_ATP_ experiments, one cell was considered one replicate. For dye-loading, to assess the effect a mutation would have on the protein, the difference in dye loading evoked by 70 mmHg PCO_2_ or 0 Ca^2+^ conditions (from the 20 mmHg baseline) for each construct were compared, using the median value for each condition from each transfection. The Mann-Whitney U one-sided test was used for the comparison on the basis that *a priori* we expected mutations to either reduce CO_2_ sensitivity or have no effect. To compare the fEPSP size across different conditions, the non-parametric Friedman two-way ANOVA was used. Statistical analysis was performed using the python language and SciPy library.

### Reagents used

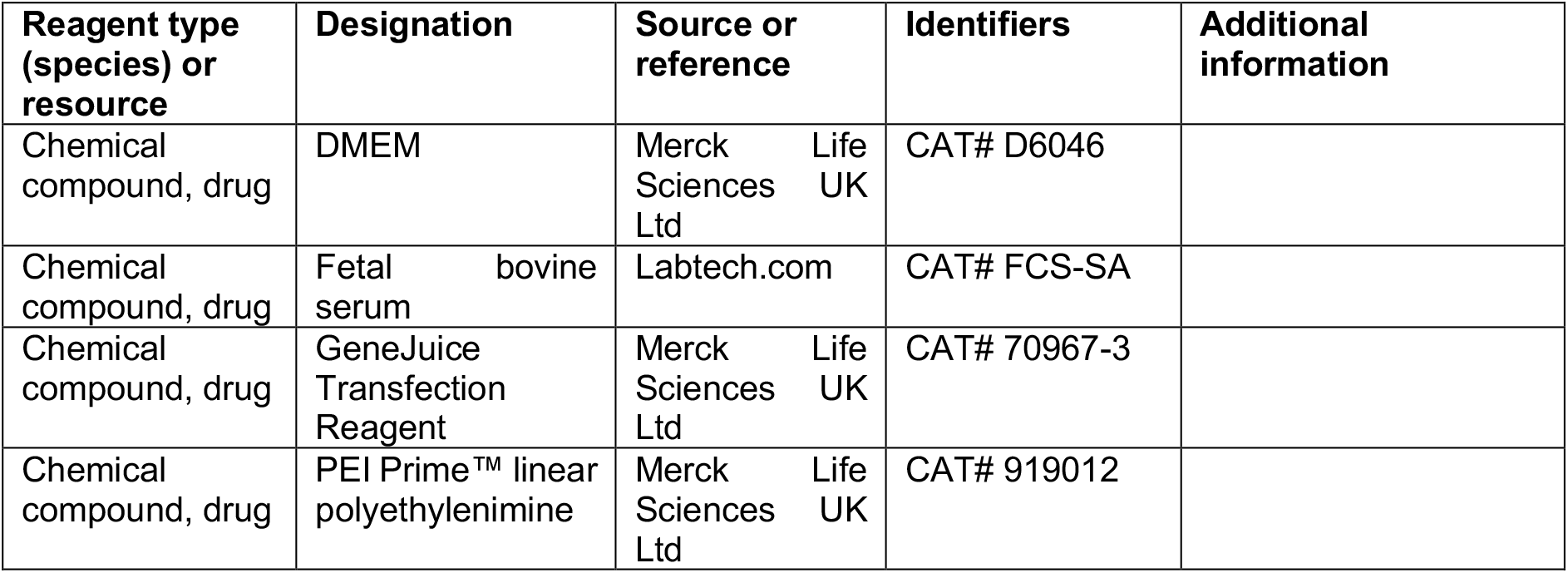
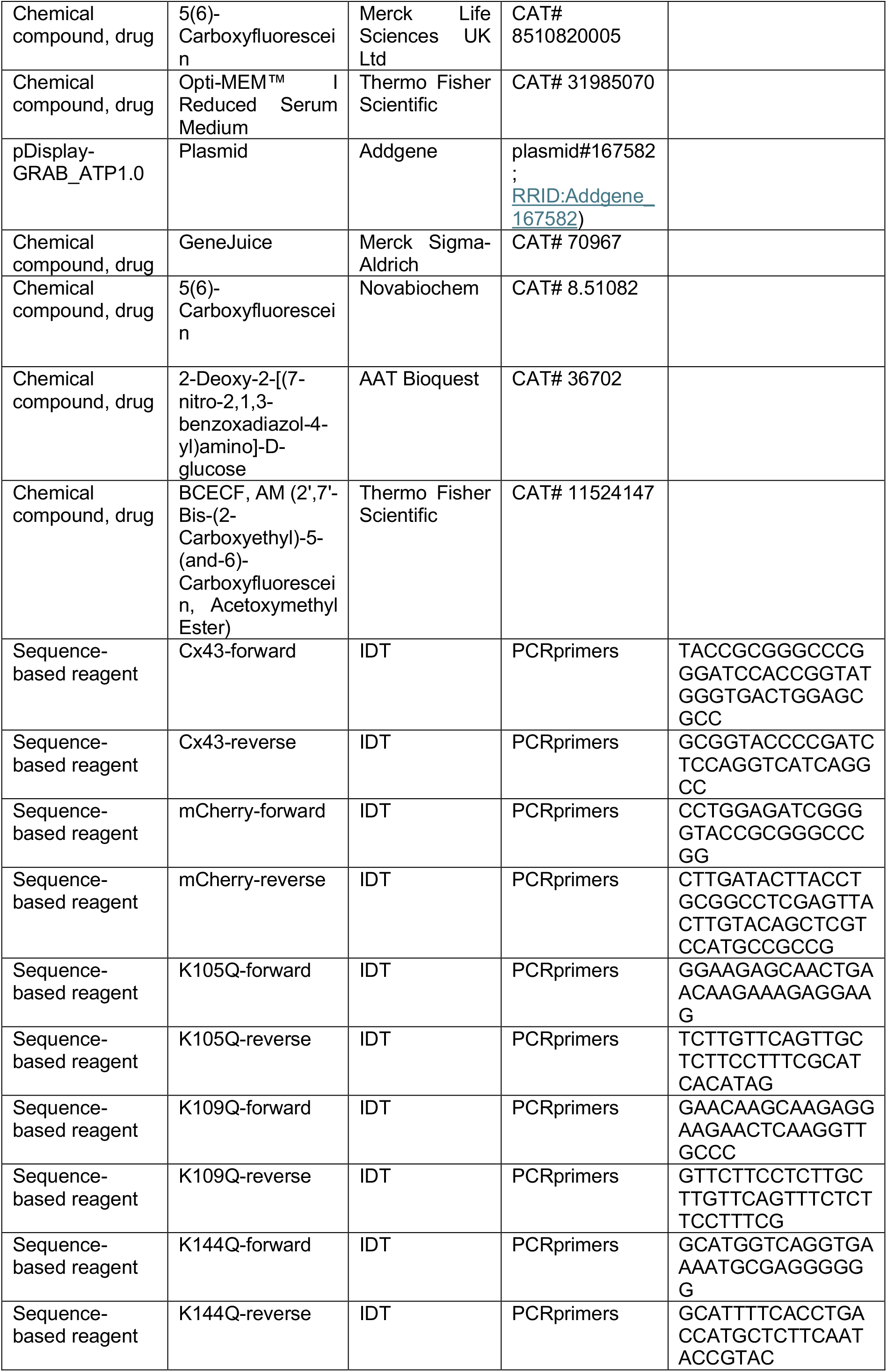
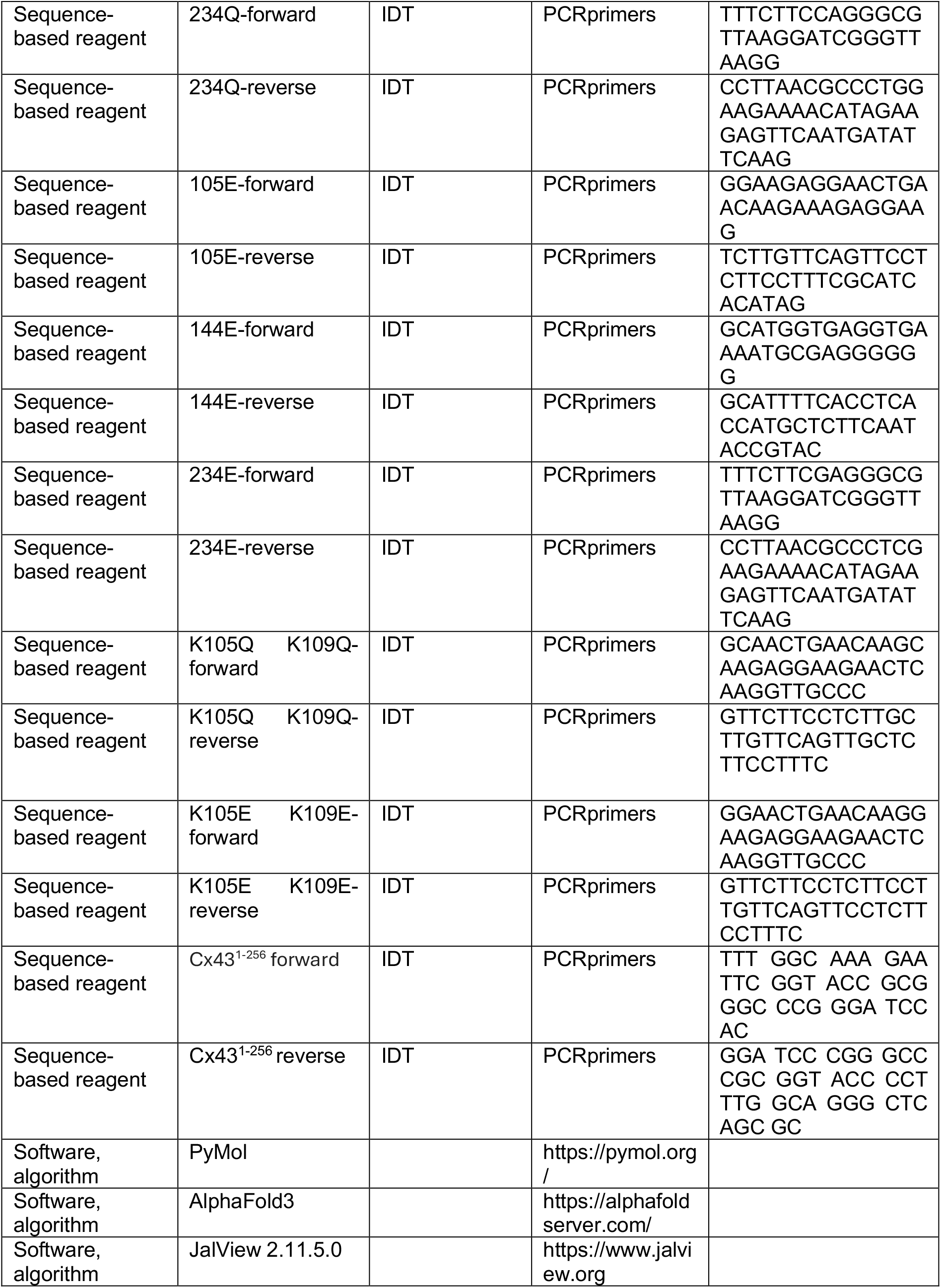

## Data availability

All data generated in the paper is available as supplements to the figures.

## Acknowledgements

We thank Prof Alexander Cameron for commenting on a draft of this paper.

---

**Figure 2, figure supplement 1:**
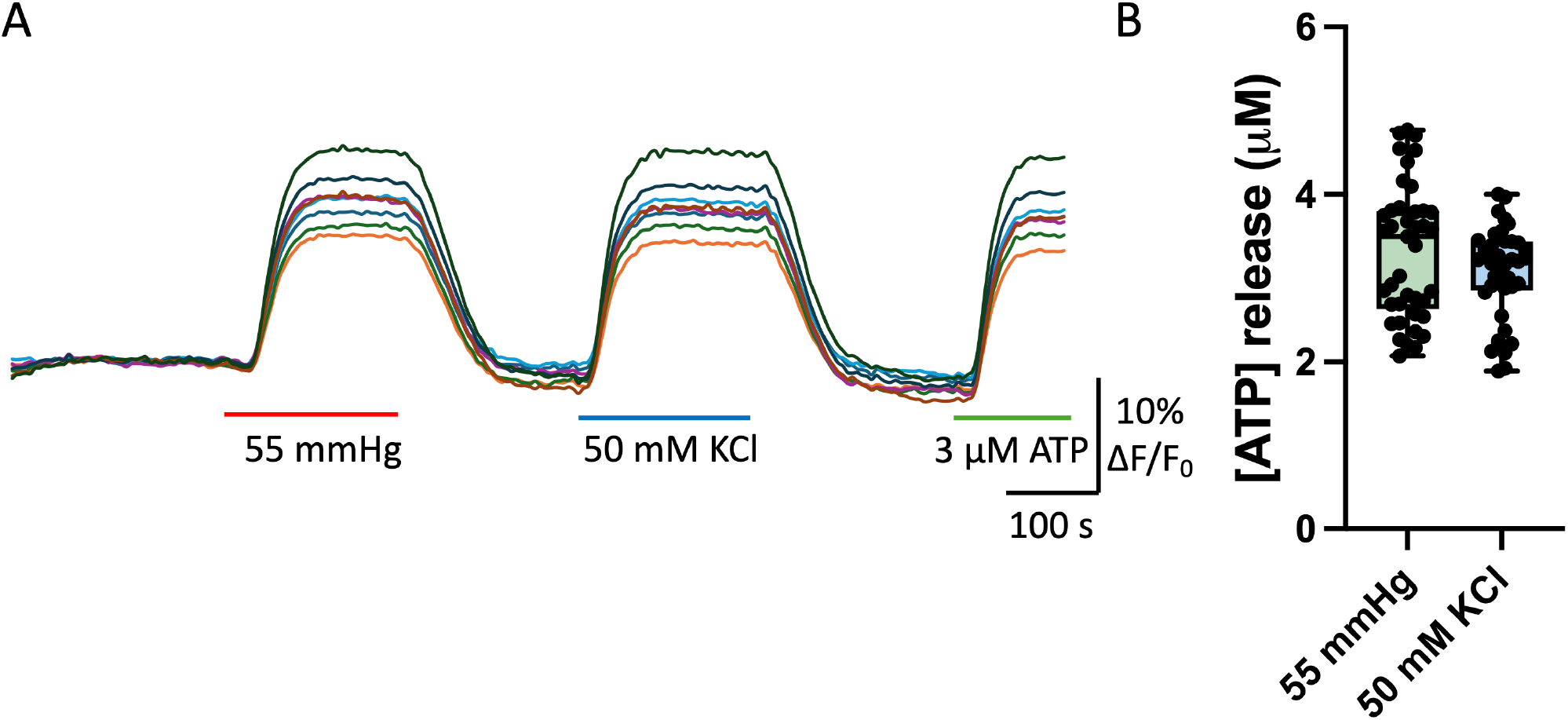
Cx43 hemichannels with a truncated C-terminus (Cx43^1-256^) retain their sensitivity to CO_2_ and depolarisation. **(A)** Changes in GRAB_ATP_ fluorescence induced by changing PCO_2_ from 20 to 55 mmHg or KCl from 3 to 50 mM. **(B)** Summary graph showing the concentration of ATP release in response to 55 mmHg PCO_2_ and 50 mM KCl, n=41 cells from 3 independent transfections.

**Figure 2, figure supplement 2:**
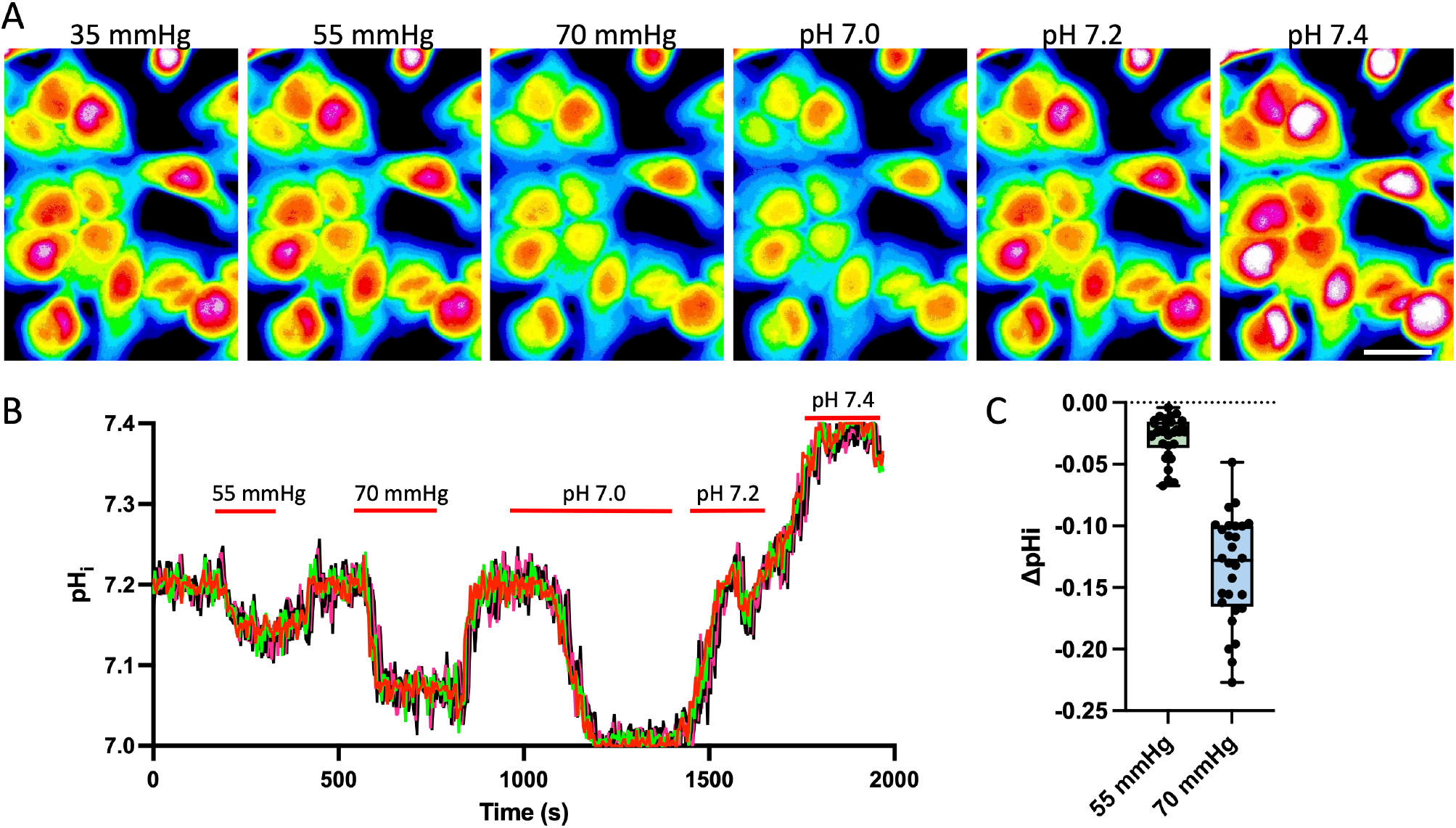
Effect of hypercapnic solutions on intracellular pH (pH_i_) of parental HeLa cells measured by BCECF fluorescence. **(A)** Images showing BCECF fluorescence at 35, 55 and 70 mmHg, and then calibration to pH 7.0, 7.2 and 7.4 of the same cells after treatment with nigericin. **(B)** Quantification of the changes in fluorescence. C) Summary plot of the change in pH_i_ induced by 55 and 70 mmHg PCO_2_. 3 independent seeds of cells, n=30.

**Figure 2, figure supplement 3:**
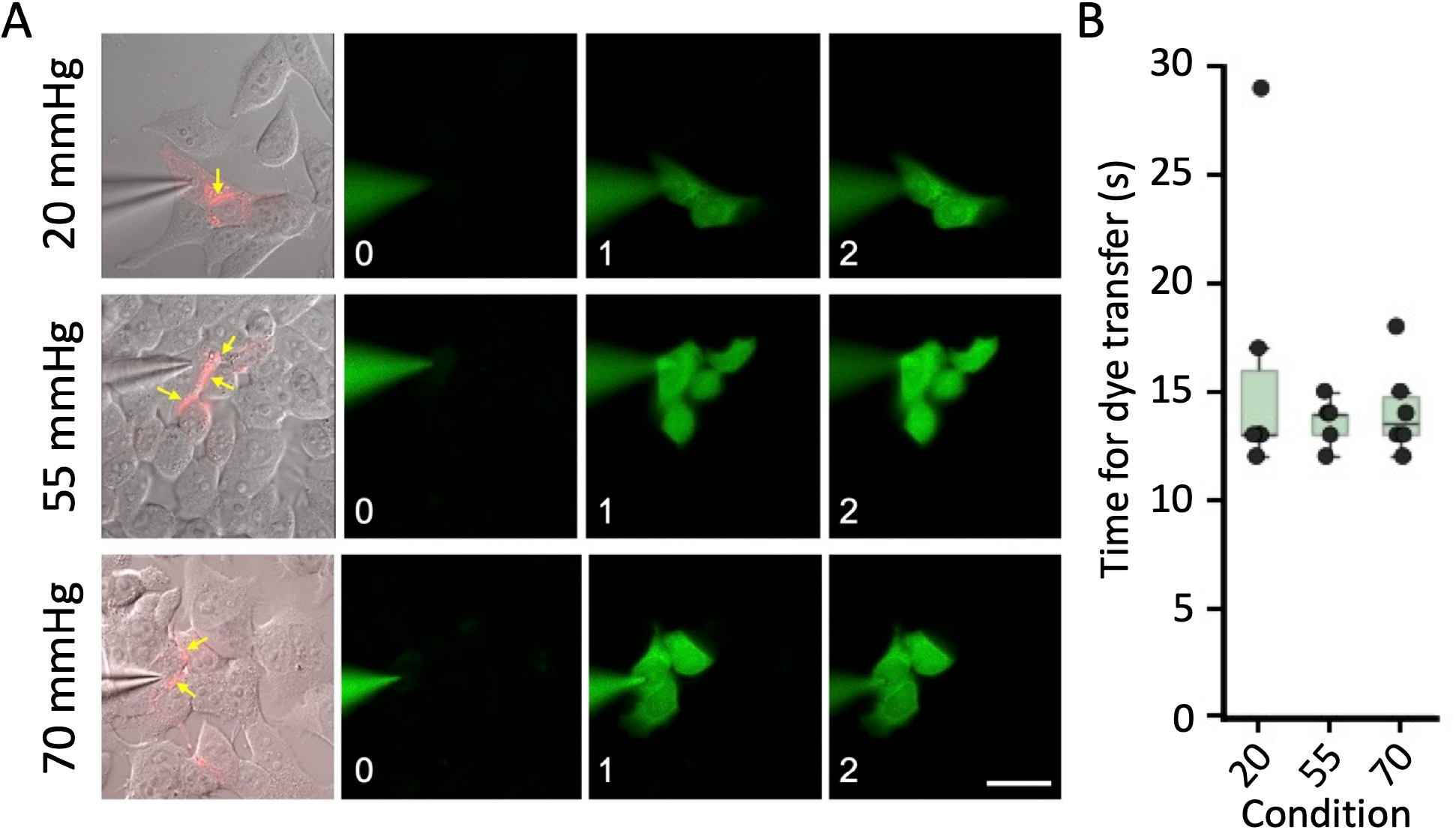
Cx43 gap junction channels are insensitive to changes in PCO_2_. **(A)** DIC image with mCherry fluorescence superimposed. Gap junctions are evident as red stripes between cells (yellow arrows). The fluorescence images show the patch pipette filled with NBDG its loading into the recorded cell and the neighbouring cells coupled via gap junctions. The numbers in each bottom left corner are the time of the recording from whole cell breakthrough in minutes. Scale bar 20 µm. **(B)** Summary graphs showing the time for the acceptor cell to reach 10% of the fluorescence of the donor cell at three different levels of PCO_2_ (n=6 for each condition).

**Figure 3, figure supplement 1:**
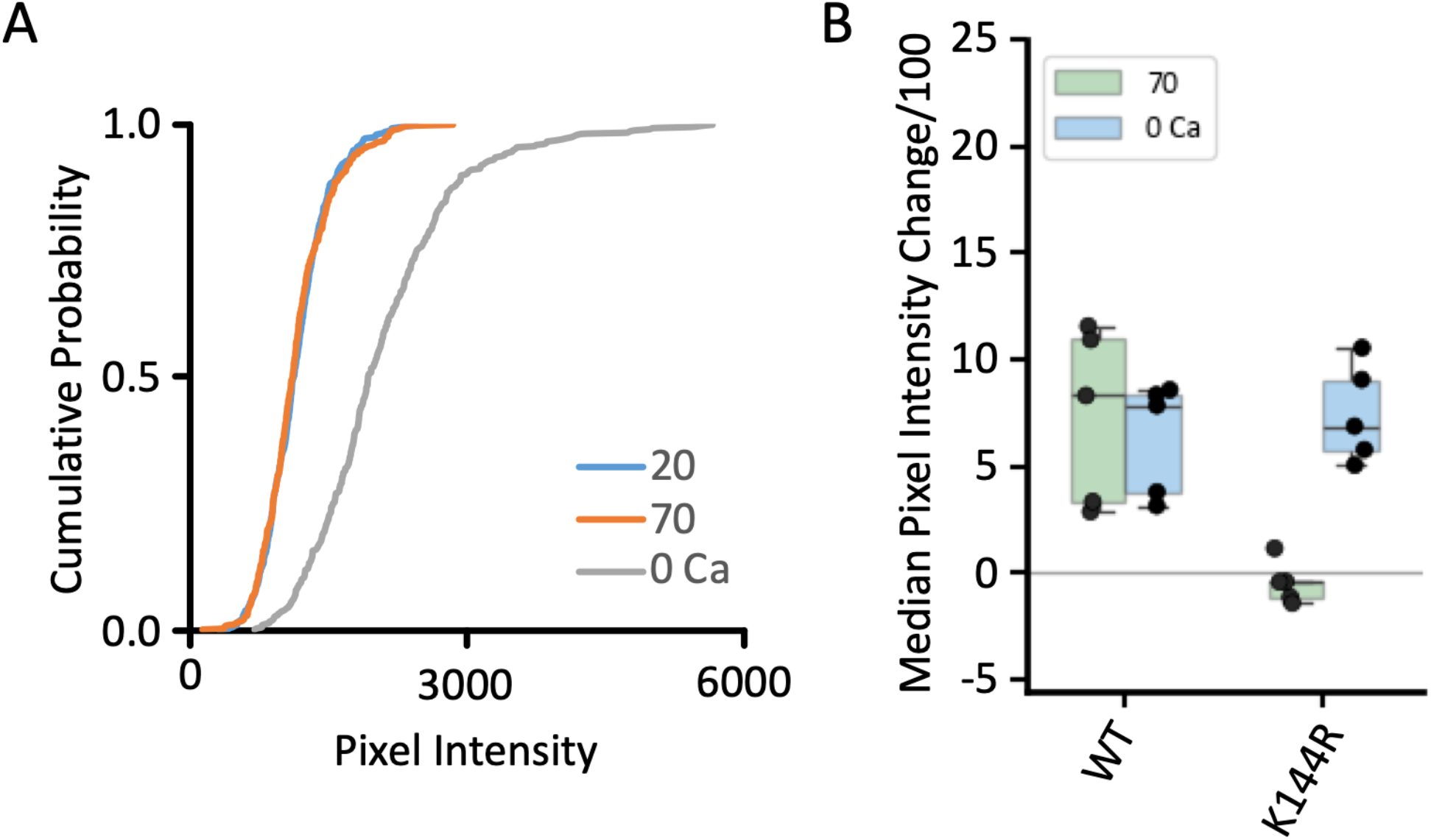
Cx43^K144R^ hemichannels are insensitive to changes in PCO_2_. **(A)** Cumulative probability plot from 5 independent transfections showing dye loading in response to 20 nd 70 mmHg and the zero Ca^2+^ stimulus. **(B)** Summary graph showing the change in median fluorescence intensity caused by 70 mmHg and zero Ca^2+^ in Cx43^WT^ (data recalculated from Fig 2) and Cx43^K144R^.

**Figure 4, figure supplement 1:**
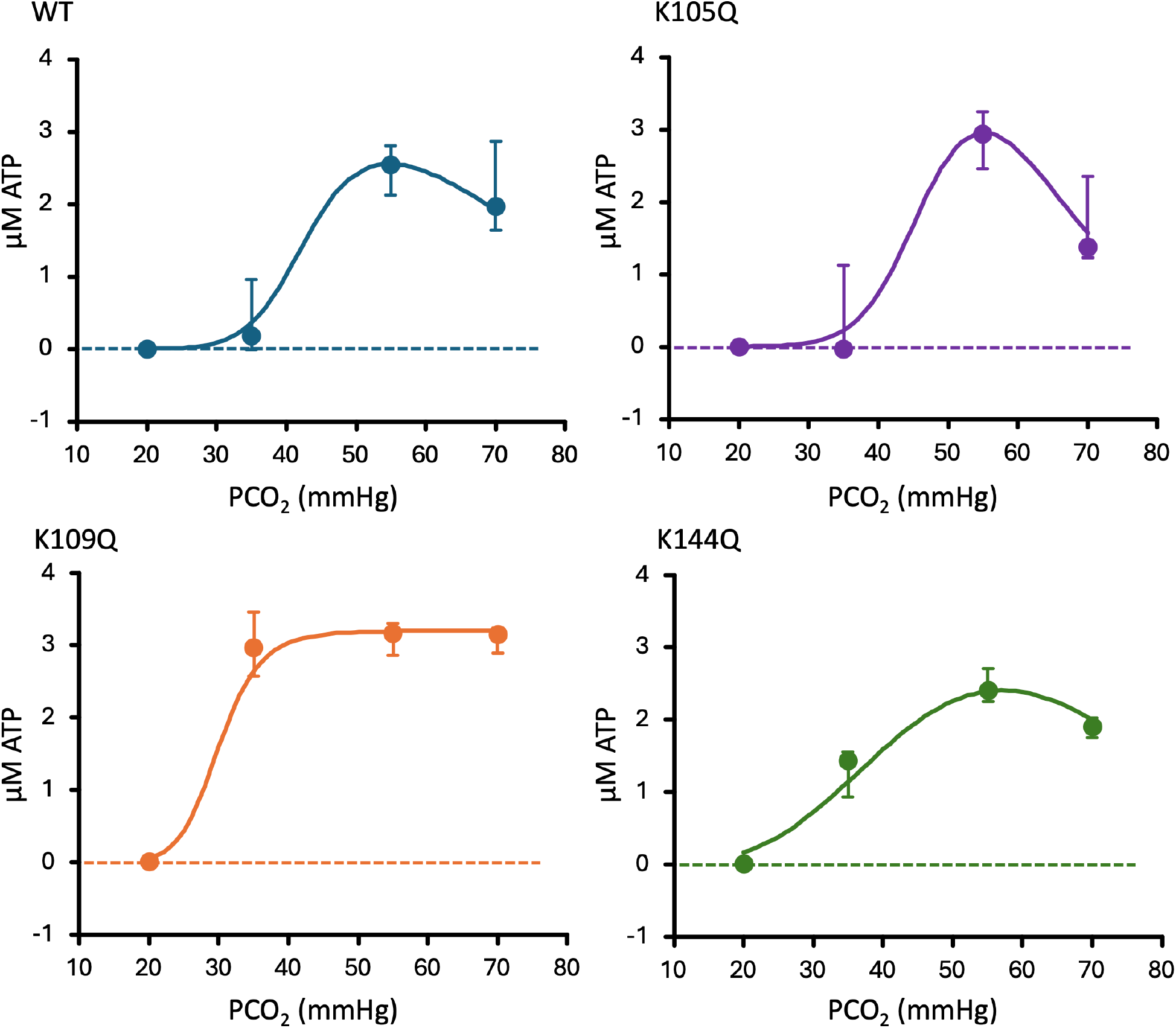
The effect of single Lys mutations on the CO_2_ dose-response properties of Cx43. Comparison of Cx43^WT^ (data replotted from Fig 2C), Cx43^K105Q^, Cx43^K109Q^ (data replotted from Fig 4E) and Cx43^K144Q^ (data replotted from Fig 4F). Data is plotted as medians with lower and upper quartiles. For the WT, K105Q and K144Q the fitted curve is drawn according to a modified Hill equation: [ATP]=Max_ATP_.[(PCO_2_/K)^H^/(1+(PCO_2_/K)^H^)].[1−(PCO_2_/K_i_)^Hi^/(1+(PCO_2_/K_i_)^Hi^)] Where K and H are the affinity and Hill coefficient of the channel for opening by CO_2_, K_i_ and H_i_ are the affinity and Hill coefficient for inhibition of the channel by CO_2_, and Max_ATP_ is the asymptotically maximum release of ATP to CO_2_. For K109Q which did not exhibit CO_2_-dependent inhibition at high levels of PCO_2_, the Hill equation was used. The parameters for the curves are:

